# APP and β-amyloid modulate protein aggregation and dissociation from recycling endosomal and exosomal membranes

**DOI:** 10.1101/2024.03.28.586966

**Authors:** Preman J. Singh, Bhavna Verma, Adam Wells, Cláudia C. Mendes, Dali Dunn, Ying-Ni Chen, Jade Oh, Lewis Blincowe, S. Mark Wainwright, Roman Fischer, Shih-Jung Fan, Adrian L. Harris, Deborah C. I. Goberdhan, Clive Wilson

**Affiliations:** Department of Physiology Anatomy and Genetics, University of Oxford, Oxford, UK; US Department of Veterans Affairs, Veterans Affairs Medical Center, Fayetteville, AR, USA; Department of Life Sciences, National Central University, Taoyuan City, Taiwan; Target Discovery Institute, University of Oxford, Oxford, UK; Department of Oncology, University of Oxford, Oxford, UK; Nuffield Department of Women’s and Reproductive Health, University of Oxford, Oxford, UK

**Author notes:** contributed equally (Preman J. Singh, Bhavna Verma). contributed equally (Deborah C. I. Goberdhan, Clive Wilson).

**Keywords:** Alzheimer’s Disease, Transforming Growth Factor-Beta-Induced, dense-core granules, secretory vesicles, endolysosomal trafficking, Rab11, corneal dystrophy, exosomes, *Drosophila melanogaster*, secondary cell

## Abstract

Secretory proteins frequently aggregate into non-soluble dense-core granules (DCGs) in recycling endosome-like compartments prior to release. By contrast, aberrantly processed Aβ-peptides derived from Amyloid Precursor Protein (APP) form pathological amyloidogenic aggregations in late-stage Alzheimer’s Disease (AD) after secretion. By examining living *Drosophila* prostate-like secondary cells, we show both APP and Aβ-peptides affect normal DCG biogenesis. These cells generate DCGs and secreted nanovesicles called Rab11-exosomes within enlarged recycling endosomes. The fly APP homologue, APP-like (APPL), associates with Rab11-exosomes and the compartmental limiting membrane, from where its extracellular domain controls protein aggregation. Proteolytic release of this membrane-associated domain permits aggregates to coalesce into a large central DCG. Mutant Aβ-peptide expression, like *Appl* loss-of-function, disrupts this assembly step and compartment motility, and increases lysosomal targeting, mirroring pathological events reported in early-stage AD. Our data therefore reveal a physiological role for APP in membrane-dependent protein aggregation, which when disrupted, rapidly triggers AD-relevant intracellular pathologies.

## Introduction

Alzheimer’s Disease (AD) is a progressive neurodegenerative disorder, which affects an increasing proportion of the world’s ageing population (Gustavsson et al., 2022). In post-mortem brains, a common feature of AD is accumulation of extracellular amyloid plaques, primarily containing organised β-strand assemblies of Aβ-peptides, which are generated from aberrant cleavage of the transmembrane Amyloid Precursor Protein (APP; Thal et al., 2002; Acquasaliente et al., 2022; Weglinski et al., 2023). The levels of Aβ-peptides and other extracellular APP cleavage products formed by neurons are determined by the activities of at least three proteases, α-, β- and γ-secretase, with β- and γ-secretase required to generate amyloidogenic Aβ-peptides (O’Brien et al., 2011; Zhang et al., 2019). Indeed, putative activating mutations in Presenilin 1, a subunit of the intramembrane γ-secretase, are, in fact, linked to rare inherited forms of AD (Kabir et al., 2020).

Although late-stage amyloid plaque pathology can compromise brain function, plaque formation is probably not a key player in the initiation of neurodegeneration in AD (Selkoe and Hardy, 2016). Furthermore, recently developed therapies that use antibodies to disrupt amyloid plaques appear to slow disease progression, but neither block nor reverse it (Travis, 2023). One hypothesis to explain the initiation of AD pathology is that AD-associated defects in the secretory pathway that releases cleaved forms of APP, including Aβ-peptides, have direct detrimental effects on neuronal cell biology, such as altered endolysosomal trafficking, which progressively leads to cell death (Kimura and Yanagisawa, 2018; Cataldo et al., 2000).

Some studies have suggested that Aβ-peptides can induce *APP* loss-of-function phenotypes, raising the possibility that these peptides might mediate their early pathological effects by interfering with normal neuronal APP functions (Bignante et al., 2013; Kepp, 2016). APP is proposed to have multiple physiological roles, for example regulating brain development, memory and synaptic functions (Nalivaeva and Turner, 2013), as well as displaying potential trophic activities (Dawkins and Small, 2014). At a molecular level, APP can act as a Wnt receptor (Liu et al., 2021) and may signal through interaction with heterotrimeric G-proteins (Copenhaver and Kögel, 2017). However, it seems unlikely that such signalling explains the full range of reported physiological functions. Nor does it indicate what the normal function, if any, of APP cleavage in secretory compartments might be.

Humans express three APP-like molecules, APP, APP-like protein 1 (APLP1) and APLP2. By contrast, the fruit fly, *Drosophila melanogaster*, has a single APP homologue, APP-like, APPL, facilitating loss-of-function studies. Despite limited conservation between sequences within the Aβ-peptide region of APPL, the protein contains α-, β- and γ-secretase cleavage sites, which also generate extracellular APP cleavage products, including Aβ-like peptides that can induce neurodegeneration (Carmine-Simmen et al., 2009; Wentzell and Kretzschmar, 2010). In addition, *Appl* loss-of-function phenotypes can be rescued by human APP, suggesting functional conservation (Luo et al., 1992). Although the specific molecular and subcellular functions of APPL remain unclear, the protein has been reported to control the balance between regulated secretion and retrograde transport at the synapse (Penserga et al., 2019).

Regulated secretion of proteins from neurons and glands typically involves the formation of secretory compartments in which proteins are condensed into non-soluble dense-core granules (DCGs), which dissipate upon extracellular release (Gondré-Lewis et al., 2012). For some signals, including certain pituitary hormones, this protein aggregation event also appears to involve amyloid formation (Maji et al., 2009). Some of the proteins regulating DCG compartment biogenesis within the *trans*-Golgi network (TGN) have been defined, and include the monomeric G-protein Arf1 (Stamnes and Rothman, 1993; Traub et al., 1993) and the adaptor protein complex, AP-1 (Bonnemaison et al., 2013). Members of the Rab family of monomeric G-proteins, which control compartment identity and vesicle trafficking, are also involved. Rab6 coats compartments emerging from the TGN (Miserey-Lenkei et al., 2010), while mature DCG compartments often interact with the recycling endosomal marker Rab11 (Sugawara et al., 2009).

We have analysed DCG biogenesis in the prostate-like secondary cells (SCs) of the male accessory gland (AG) in the fruit fly, *Drosophila melanogaster* (Figure 1A). They form DCG compartments that are thousands of times larger in volume than other secretory cells (Redhai et al., 2016), allowing intra-compartmental events to be analysed by fluorescence microscopy. SC DCG compartments are marked by Rab11 and also generate intraluminal vesicles (ILVs), which are secreted as exosomes (Corrigan et al., 2014; Fan et al., 2020). Since these exosomes are not generated in late endosomes, the previously identified source of exosomes, and they carry distinct cargos, they are termed Rab11-exosomes (Fan et al., 2020; Marie et al., 2023).

**Figure 1.**
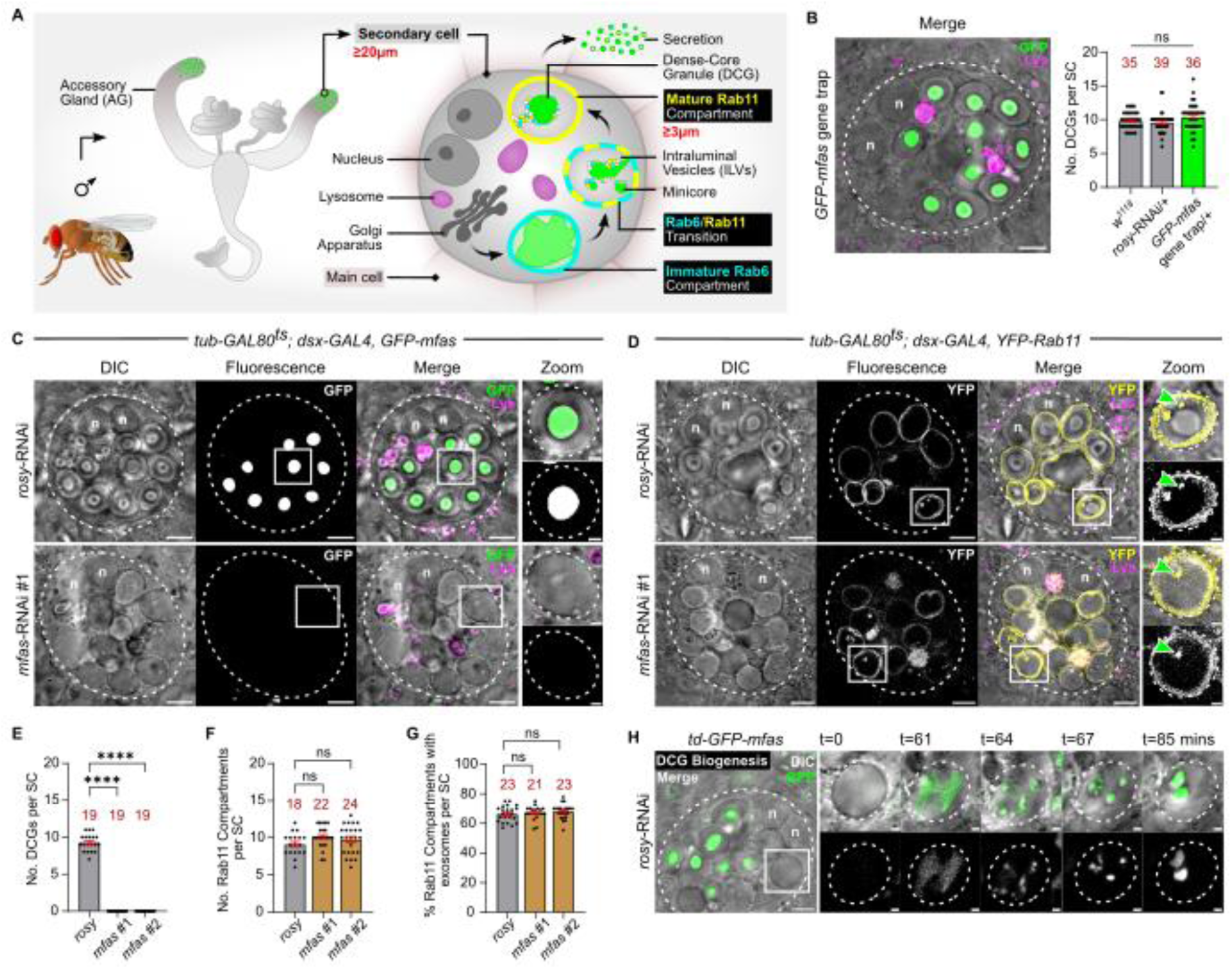
*Drosophila* TGFBI selectively drives DCG assembly in SCs. (A) Schematic showing male accessory gland, secondary cells (SCs) at the distal tip, and key compartments within the SC. Note that mature DCG compartments containing ILVs, which are the precursors of Rab11-exosomes, are generated following a Rab6 to Rab11 transition. (B) *Ex vivo*, wide-field fluorescence micrograph merged with differential interference contrast (DIC) image of SC from 6-day-old male expressing *GFP-mfas* gene trap, showing fusion protein concentrated inside DCGs within DCG compartments. *GFP-mfas* does not affect the number of DCGs per SC, when compared to *w^1118^* and *UAS-rosy-*RNAi controls. (C) *Ex vivo*, wide-field fluorescence micrographs and DIC images of SCs from 6-day-old males expressing *GFP-mfas* gene trap and SC-specific *rosy*-RNAi or *mfas*-RNAi. Note in *mfas* knockdown that large compartments lack DCGs, which are normally readily identified by DIC (shown at high magnification for compartments outlined with white boxes in Zoom panels). (D) *Ex vivo*, wide-field fluorescence micrographs and DIC images of SCs from 6-day-old males expressing *YFP-Rab11* and SC-specific *rosy*-RNAi or *mfas*-RNAi. Note that similar to controls, large empty compartments are Rab11-positive and the majority contain Rab11-positive ILVs (green arrowhead in Zoom). (E-G) Bar charts showing that knockdown of *mfas* with two independent RNAis completely blocks DCG biogenesis (E), but has no effect on numbers of Rab11-positive large compartments (F) or the proportion of these compartments containing Rab11-positive ILVs (G). (H) Stills from time-lapse movie of DCG biogenesis in SC from 6-day-old male expressing *GFP-mfas* gene trap, focusing on compartment marked by white box (DIC and GFP in top row; GFP in bottom row). In this example, multiple mini-cores form from a GFP-MFAS cloud, then they coalesce. *td-GFP-mfas* = *tub-GAL80^ts^/+; dsx- GAL4, GFP-mfas/+*. In all images, approximate cell boundary and compartment boundaries are marked with dashed white line; n = nuclei of binucleate cells; LysoTracker Red (magenta) marks acidic compartments in B-D. Scale bars = 5 µm and 1 µm in Zoom. For bar charts, data are represented as mean ± SEM and were analysed using the Kruskal-Wallis test, followed by Dunn’s multiple comparisons post hoc test; n = number above bar, ****P<0.0001, ns = not significant. See also Figure S1, and Movies S1 and S2.

Recently, we showed that SC DCG biogenesis requires Arf1 and AP-1, and using real time imaging, demonstrated that Rab6-labelled DCG precursor compartments must receive a Rab11-mediated input to initiate a Rab6 to Rab11 transition that rapidly triggers Rab11-exosome and DCG biogenesis (Figure 1A; Wells et al., 2023; Loh et al., 2023). The ILVs produced surround the DCG and form chains that link the DCG to the compartment’s limiting membrane (Figure 1A; Fan et al., 2020). These ILVs appear to be involved in normal DCG formation (Marie et al., 2023; Dar et al., 2021). Here, we further investigate the genetic regulation of SC DCG biogenesis. We show that DCG protein aggregation is controlled by the homologues of two human proteins involved in amyloidogenic diseases. First, Midline Fasciclin (MFAS), the *Drosophila* homologue of Transforming Growth Factor-β-Induced (TGFBI), a protein mutated in many forms of dominant amyloid corneal dystrophy (Han et al., 2016), is essential for aggregation. Second, APPL regulates the membrane- and ILV-dependent priming of TGFBI aggregation, and when proteolytically cleaved, permits aggregated TGFBI to coalesce and mature into a single large DCG. Furthermore, expressing pathological human Aβ-peptide mutants aborts this maturation process, inhibits normal compartment motility and leads to increased endolysosomal targeting of secretory compartments, mirroring some of the earliest defects observed in AD and suggesting a link between APP’s secretory functions and AD pathology.

## Results

### *Drosophila* TGFBI drives DCG assembly in SCs

We screened publicly available gene trap lines (Nagarkar-Jaiswal et al., 2015), where green fluorescent protein (GFP) fusion proteins are expressed from the endogenous locus of specific genes, for genes transcribed at high levels in the AG (Leader et al., 2018). One gene, *mfas* (Hu et al., 1998), which encodes the homologue of the secreted fibrillar protein TGFBI (Nielsen et al., 2020; Corona and Blobe, 2021), was highly expressed specifically in SCs (Figure 1B), a result consistent with recent single-cell RNAseq analysis (Immarigeon et al., 2021; Li et al., 2022). This protein is expressed in many human tissues. However, dominant mutations in TGFBI selectively affect the cornea, leading to most cases of corneal dystrophy, where amyloid deposits progressively restrict vision (Han et al., 2016). GFP-MFAS was concentrated in all DCGs of SCs; in heterozygous gene trap flies, it did not affect the overall morphology of SCs or their DCGs, which can be visualised independently by differential interference contrast microscopy (DIC; Figure 1B and 1C). SCs in males heterozygous for the gene trap and carrying transgenes that permit adult SC-specific expression of other genes via the GAL4/UAS system under temperature-sensitive GAL80^ts^-inducible control (Fan et al., 2020), also produced normal numbers of DCG compartments (Figure S1D).

Knockdown of *mfas* with two independent RNAis specifically in adult SCs produced large secretory compartments with no DCGs (Figures 1C, 1E and S1A). Normally, all DCG compartments in SCs are marked by Rab11 (Fan et al., 2020), which can be visualised using a *YFP-Rab11* protein fusion expressed from the endogenous gene locus (Figure 1D; Dunst et al., 2015). The number of compartments marked by Rab11 was unaltered following *mfas* knockdown when compared to control cells expressing an RNAi targeting the xanthine dehydrogenase gene, *rosy*, which did not affect DCG biogenesis (Figures 1D, 1F and S1B; Marie et al., 2023). Furthermore, the number of compartments labelled with Rab6, which marks precursor compartments that lack DCGs as well as less mature DCG compartments, was also unaffected (Figures S1C and S1E; Wells et al., 2023). Rab11 and Rab6 label a fraction of the ILVs present in DCG compartments (Fan et al., 2020; Wells et al., 2023). By counting the proportion of labelled compartments containing Rab11- and Rab6-positive puncta in their lumen, we concluded that there was no major effect on ILV biogenesis in these compartments (Figures 1D, 1G, S1B, S1C and S1F). Therefore, MFAS, the homologue of a human protein with amyloidogenic potential, has a highly selective, essential role in normal protein aggregation that drives large DCG formation in SCs.

Using the *mfas* gene trap line, DCG biogenesis was followed in real-time. We observed two aggregation processes. In the first, multiple dispersed mini-cores initially assembled by rapid condensation of diffuse clouds of GFP-MFAS present within immature compartments (Figure 1H; Movie S1). These mini-cores then mobilised inside the compartment and fused with each other to make a large DCG within approximately 30 minutes. Alternatively, in about half of maturing compartments, the entire DCG condensed in minutes from a large diffuse central GFP-MFAS cloud; individual mini-cores were less readily identifiable, though the aggregation events were not entirely uniform across the cloud (Figure S1G; Movie S2). Overall, our data suggest that MFAS drives the protein aggregation required for SC DCG biogenesis by eliciting rapid condensation events in maturing compartments.

### GAPDH is required for mini-core fusion, and co-isolates with human Rab11-exosomes and other AD-associated glycolytic enzyme biomarkers

The mini-core phase observed in some examples of DCG biogenesis (Figure 1H) was reminiscent of a phenotype previously observed after knockdown of the *Drosophila* glyceraldehyde 3-phosphate dehydrogenase isoform, *GAPDH2*, in SCs, where multiple mini-cores are produced in each compartment (Dar et al., 2021). Knockdown of this glycolytic enzyme in SCs expressing the *mfas* gene trap led to the production of a slightly increased number of DCG compartments compared to *rosy*-RNAi-expressing controls. Almost all of these compartments contained multiple mini-cores, many in close proximity to the compartment’s limiting membrane (Figure 2A-C).

**Figure 2.**
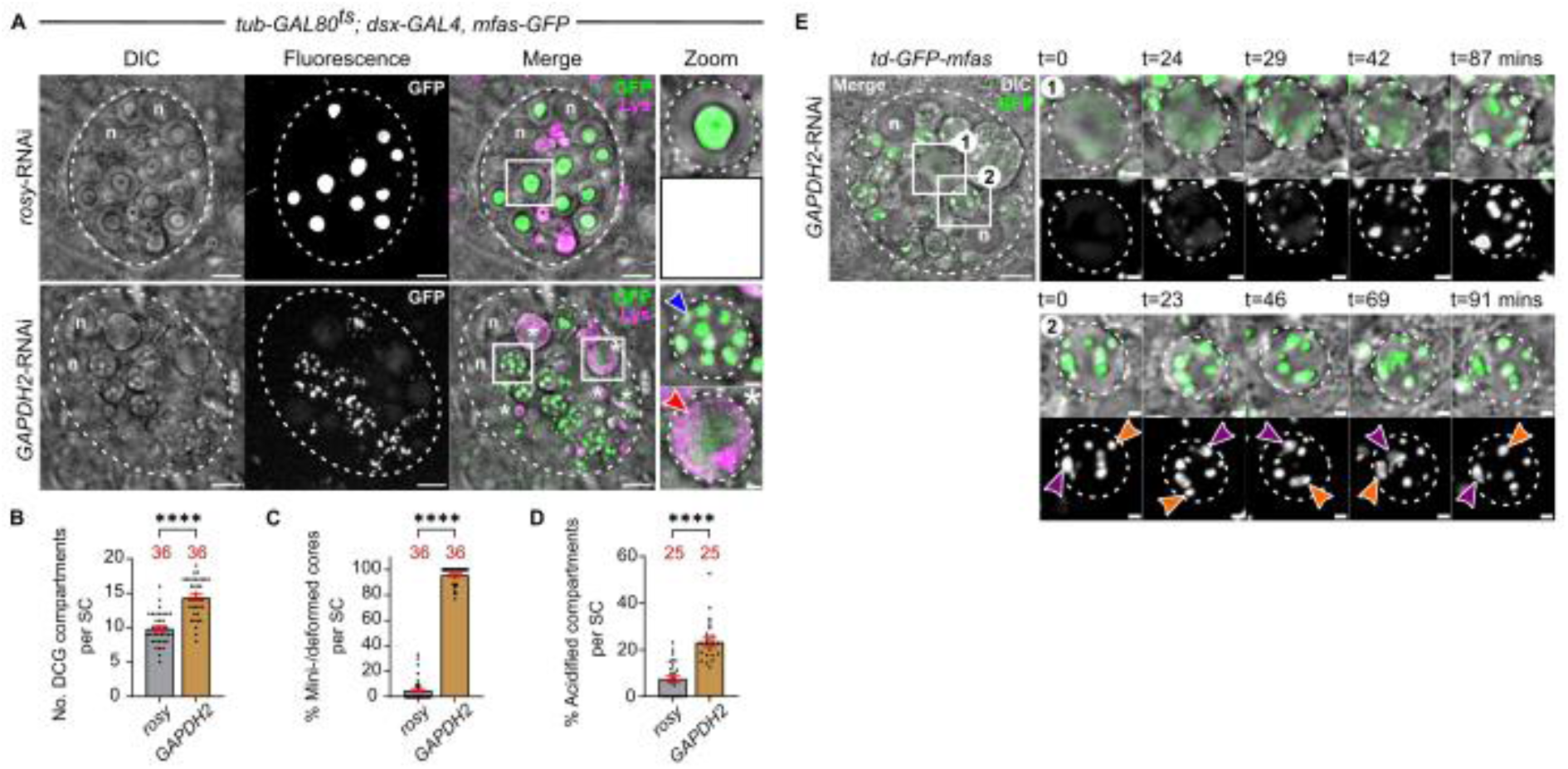
GAPDH2 is required for mini-core fusion in SCs. (A) *Ex vivo*, wide-field fluorescence micrographs and DIC images of SCs from 6-day-old males expressing *GFP-mfas* gene trap and SC-specific *rosy*-RNAi or *GAPDH2*-RNAi. Following knockdown of *GAPDH2* note that large secretory compartments contain multiple mini-cores (blue arrowhead), labelled with GFP-MFAS and visible by DIC (shown for compartments outlined with white boxes in Zoom panels). White asterisks and white boxes marked with asterisks in the Merge panel indicate DCG compartments with acidification phenotype (lower zoom panel in *GAPDH2* knockdown; red arrowhead marks acidic domain). (B-D) Bar charts showing that knockdown of *GAPDH2* slightly increases the number of DCG compartments (B). Almost all of these compartments contain multiple mini-cores (C). A greater proportion of large compartments display the DCG acidification phenotype than in controls (D). (E) Stills from time-lapse movie of DCG biogenesis in SC from 6-day-old male expressing *GFP-mfas* gene trap and *GAPDH2*-RNAi, focusing on an immature DCG compartment (top rows [1]) and a mature compartment (bottom rows [2]), both marked by white boxes. Multiple mini-cores condense from clouds of GFP-MFAS during maturation and remain motile within compartments that also rotate within the cell (two mini-cores marked by purple and orange arrowheads in [2]). However, unlike control cells (Figure 1H), even if these mini-cores collide, they do not appear to fuse. In all images, approximate cell boundary and compartment boundaries are marked with dashed white line; n = nuclei of binucleate cells; LysoTracker Red (magenta) marks acidic compartments in A. Scale bars = 5 µm and 1 µm in Zoom. For bar charts, data are represented as mean ± SEM and were analysed using the Mann-Whitney test; n = number above bar, ****P<0.0001, ns = not significant. See also Figure S2, Movie S3 and Table S1.

We previously showed that GAPDH plays a conserved, so-called ‘moonlighting’, role in humans and flies in clustering Rab11-exosomes and other secreted extracellular vesicles (EVs; Dar et al., 2021). In SCs, DCGs are partially coated with strings of ILVs that extend in chains to the limiting membrane of the DCG compartment (Figure 1A; Fan et al., 2020; Wells et al., 2023). Following *GAPDH2* knockdown, most mini-cores formed are located near the compartment boundary and are frequently surrounded by ILVs, which are therefore also located peripherally (Dar et al., 2021). We hypothesised that the reduced ILV clustering activity in GAPDH2 knockdown cells might either inhibit mini-core movement towards the compartment centre or suppress fusion during DCG maturation. Indeed, time-lapse imaging revealed that the mini-cores in SCs with *GAPDH2* knockdown still formed and both these mini-cores and the DCG compartments that contained them were mobile, similar to normal cells. However, the mini-cores often failed to collide and when they seemed to make contact, no fusion took place (Figure 2E; Movie S3).

We also observed that *GAPDH2* knockdown SCs contained several large compartments with diffuse or no GFP-MFAS that were contacted by one or more peripheral crescent-shaped acidic structures, stained by the acid-sensitive dye LysoTracker Red (Figure 2A). Real-time imaging revealed that this phenotype only appeared occasionally in control cells (Figure 2D), but was generated by the fusion of DCG compartments with lysosomal structures and subsequent rapid dispersion of the DCG (Figure S1G). We therefore refer to this phenotype as a ‘DCG acidification phenotype’, and conclude that it can occur more frequently in cells where DCG biogenesis is defective.

We further investigated whether there is an evolutionarily conserved association between Rab11-exosomes and GAPDH. In humans, exosomes are formed as ILVs in Rab11a-labelled, recycling endosomal compartments in several human cancer cell lines (Fan et al., 2020). Previous work has shown that levels of the accessory ESCRT-III proteins, which play conserved and selective roles in Rab11-exosome biogenesis, are increased in small EV (sEV) preparations from human cancer cells that are enriched in Rab11a-exosomes (Marie et al., 2023). To determine whether GAPDH might also be enriched in these exosomes, we compared small EV (sEV) preparations made from HCT116 human colorectal cancer cells under glutamine-depleted versus glutamine-replete conditions. Glutamine depletion reduces the activity of the nutrient-sensing kinase complex, mechanistic Target Of Rapamycin Complex 1 (mTORC1), and this promotes Rab11a-exosome secretion (Fan et al., 2020). Western blots of sEV preparations revealed that this treatment increased the levels of EV-associated GAPDH, as well as Rab11a, while levels of CD63, a late endosomal exosome marker, were reduced (Figure S2).

Interestingly, in humans, both exosomes and GAPDH are proposed to be associated with β-amyloid plaques and to play a role in their assembly (Yuyama and Igarashi, 2017; Itakura et al., 2015; Lazarev et al., 2021). Furthermore, increased circulating levels of modified forms of extracellular GAPDH are associated with Aβ-peptide burden in AD patients (Tsai et al., 2020). Levels of two other glycolytic enzymes, fructose-bisphosphate aldolase A (ALDOA) and pyruvate kinase (PKM), have recently been shown to be elevated in cerebrospinal fluid (CSF) of AD patients, together with four other hub proteins, forming an AD signature (Li et al., 2022). This meta-analysis highlighted ‘extracellular exosome’ as the most highly represented GO term for proteins enriched in CSF of AD patients. Interestingly, ALDOA and PKM were two of only 48 proteins identified as Rab11a-exosome-enriched markers in a recent comparative proteomic analysis of HCT116 sEVs (Marie et al., 2023).

We tested whether these two glycolytic enzymes might be elevated in Rab11a-exosome-enriched preparations from another cell type, the human HeLa cervical cancer cell line (Fan et al., 2020). We undertook a comparative proteomics analysis of sEVs isolated from these cells by size-exclusion chromatography, either under glutamine-replete or glutamine-depleted conditions. A total of 1156 proteins were detected in this screen of which 61 were significantly increased in Rab11a-exosome-enriched EV preparations (Table S1). Only seven proteins were enriched in both HCT116 and HeLa cell Rab11a-exosome-enriched preparations and remarkably, these included ALDOA and PKM. Furthermore, GAPDH also appeared to be increased, though the change did not reach significance (Table S1).

We conclude that specific glycolytic enzymes associated with the AD secretome are also elevated in human sEV preparations enriched in Rab11a-exosomes. In *Drosophila* SCs, one of these proteins, GAPDH, plays a role in Rab11-exosome clustering and in Rab11-exosome-regulated protein aggregation. Given these observations, the previous links between GAPDH, exosomes and β-amyloid plaques, and the proposed role of defective endosomal trafficking in the initiation of AD pathology (Cataldo et al., 2000), we hypothesised that APP might function in the Rab11-exosome trafficking pathway and this role could be misregulated in AD. We therefore tested the functions of transmembrane APP and its extracellular cleavage products (Figure 3A) in SC secretory and endosomal trafficking.

**Figure 3.**
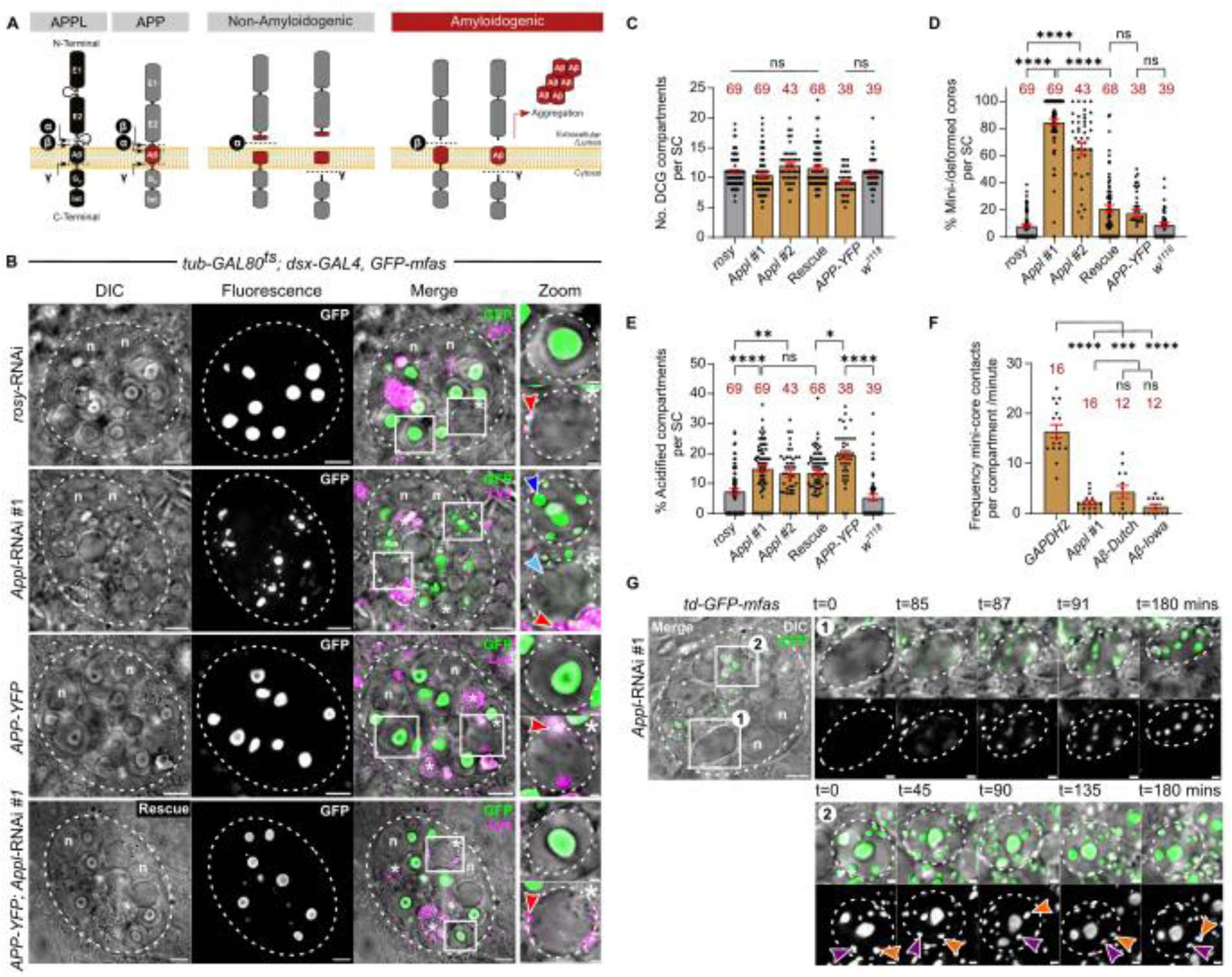
*Drosophila* APPL regulates formation of large DCGs in SCs. (A) Schematic showing structural similarities between human APP and *Drosophila* APPL proteins, as well as the location of cleavage sites for α-, β- and γ-secretases, and the APP polypeptide products that they generate. (B) *Ex vivo*, wide-field fluorescence micrographs and DIC images of SCs from 6-day-old males expressing *GFP-mfas* gene trap and SC-specific *rosy*-RNAi, *Appl*-RNAi, APP-YFP and the combination of *Appl*-RNAi and APP-YFP. Note that following knockdown of *Appl*, large secretory compartments contain multiple mini-cores (blue arrowhead), labelled with GFP-MFAS and visible by DIC (shown for compartments outlined with white boxes in upper Zoom panels). White asterisks and white boxes marked with asterisks in the Merge channel indicate DCG compartments with acidification phenotype (lower zoom panel; red arrowheads mark acidic microdomains). DCG that has not yet been dissipated marked with light blue arrowhead. (C-E) Bar charts showing that knockdown of *Appl* with two independent RNAis does not affect the number of DCG compartments (C), but most of these compartments contain multiple mini-cores, a phenotype rescued by APP-YFP co-expression with *Appl-*RNAi#1 (D). Both *Appl* knockdown and APP-YFP expression induce the DCG acidification phenotype when compared to controls (E). (F) Bar chart showing frequency of mini-core overlap in time-lapse videos over a 30-minute period for single mini-cores selected from multiple different compartments, following SC-specific *GAPDH2* and *Appl* knockdown, and expression of the human AD-associated Aβ-42 Dutch and Aβ-42 Iowa mutants. (G) Stills from time-lapse movie of DCG biogenesis in SC from 6-day-old male expressing *GFP-mfas* gene trap and *Appl*-RNAi, focusing on one compartment as it forms mini-cores (white box 1) and another mature compartment (white box 2). Note that multiple mini-cores condense from clouds of GFP-MFAS, but the compartments and mini-cores are relatively immobile (two mini-cores marked by purple and orange arrowheads in [2]) and unlike control cells (Figure 1H), they do not fuse. In all images, approximate cell boundary and compartment boundaries are marked with dashed white line; n = nuclei of binucleate cells; LysoTracker Red (magenta) marks acidic compartments in B. Scale bars = 5 µm and 1 µm in Zoom. For bar charts, data are represented as mean ± SEM and were analysed using the Kruskal-Wallis test, followed by Dunn’s multiple comparisons post hoc test; n = number above bar, *P<0.05, **P<0.01, ***P<0.001, ****P<0.0001, ns = not significant. See also Figure S3 and Movie S4.

### *Drosophila* APPL regulates formation of large DCGs in SCs

We knocked down the *Drosophila APP* homologue, *Appl*, in SCs using two independent RNAis. In both knockdowns, although the total number of large DCG compartments remained unaltered, most of these compartments contained several mini-cores, and often a mis-shapen central DCG of reduced size (Figures 3B-3D and S3A). As with *GAPDH2* knockdown, most mini-cores were in close proximity to the limiting membrane of each compartment. Furthermore, there was an increased proportion of large compartments with the DCG acidification phenotype (Figure 3B and 3E, Figure S3A).

Analysis of *Appl* knockdown cells in YFP-Rab11 and CFP-Rab6 backgrounds revealed that the Rab identity of the defective mini-core compartments and the proportion of these compartments containing Rab-labelled exosomes remained unchanged compared to controls (Figure S3B, S3C, S3I-L). However, as we previously found for *GAPDH2* knockdown (Dar et al., 2021), and unlike controls, many Rab-labelled ILVs were clustered at the periphery of each compartment near the mini-cores (Figure S3B and S3C). The compartments with a DCG acidification phenotype were not marked by either Rab11 or Rab6 (Figure S3B, S3C), consistent with them being DCG compartments that have been targeted for lysosomal degradation.

To further assess APPL’s role in DCG compartment biogenesis, we analysed SCs from males hemizygous for an *Appl* null allele, *Appl^d^* (Luo et al., 1992). *Appl* null SCs contained similar numbers of DCG compartments to controls (Figure S3D), but in contrast to *Appl* knockdown cells, mini-cores were uncommon (Figure S3A). However, DCGs in mutant cells were often misshapen and more frequently contacted the limiting membrane of the compartment (Figure S3A, S3E and S3G). Most importantly, when compared to control and *Appl* knockdown cells, *Appl* null cells contained many more acidified DCG compartments (Figure S3A and S3F). Therefore, these findings further support the conclusion that *Appl* is essential for the normal maturation of DCG compartments and suggest that APPL protein plays an important role in dissociation of DCG aggregates from the limiting membrane of these compartments.

Human APP has previously been shown to partially rescue the behavioural defects caused by *Appl* loss-of-function in flies (Luo et al., 1992). It is proteolytically cleaved by secretases in the *Drosophila* nervous system, although processing at the β-secretase site is inefficient (Greeve et al., 2004; Ramaker et al., 2016). We found that over-expression of an APP-YFP fusion protein in SCs had no significant effect on DCG number, integrity and shape when compared to controls (Figure 3A-C), but it did induce the DCG acidification phenotype (Figure 3E), perhaps because overexpressed APP is not fully cleaved in these cells. Overexpressing APP-YFP in *Appl* knockdown cells strongly rescued the mini-core phenotype, though about 10% of DCGs displayed a novel phenotype in which the core’s centre lacked fluorescent GFP-MFAS (Figure 3D and S3H). APP-YFP expression could not suppress the DCG compartment acidification associated with *Appl* knockdown, presumably because it induces this phenotype when expressed alone (Figure 3B and 3E). These findings suggest that human APP and fly APPL share the functional activity required for normal DCG protein aggregation and subsequent separation from the DCG compartment’s limiting membrane.

We investigated whether the mini-core phenotype caused by *Appl* knockdown was linked to the failure of mini-cores to collide and fuse, as we had observed for *GAPDH2* knockdown. Time-lapse movies of GFP-MFAS in *Appl* knockdown SCs revealed that compared to *GAPDH2* knockdown, the multiple mini-cores formed by GFP-MFAS condensation in each compartment rarely moved relative to each other, because the defective compartments were much more static within the cytoplasm of SCs (Figure 3F; Movie S4), suggesting that these compartments might be more stably attached to the surrounding cytoskeleton.

### Cleavage of *Drosophila* APPL accompanies normal DCG formation

Based on these findings, we hypothesised that transmembrane APPL is likely to be trafficked to DCG precursor compartments, where it interacts with aggregating proteins and is then proteolytically cleaved as part of the DCG maturation process. Reducing APPL levels or overexpressing APP that cannot be fully cleaved might disrupt these interactions, ultimately leading to lysosomal targeting of maturing compartments.

To investigate this further, an APPL construct tagged at both its N-terminal extracellular and C-terminal intracellular ends was overexpressed in SCs (dt-APPL; Figure 4A; Ramaker et al., 2016). The compartmental organisation of SCs was disrupted following six days of expression, with expanded lysosomal compartments, but it was still possible to identify compartments representing different stages of DCG biogenesis. In large DCG precursor compartments that had yet to form a DCG, the enhanced GFP- (EGFP-) labelled extracellular domain (ECD) was located on and just inside the compartment’s limiting membrane. The membrane fluorescence co-localised with the monomeric RFP- (mRFP-) marked intracellular domain (ICD; Figure 4C, box 1) at the membrane. In mature DCG compartments, all the ECD, but no ICD, was concentrated in the DCG itself. In many, but not all, compartments, low levels of the ICD, which is not detached from APPL’s transmembrane domain following cleavage by α- or β-secretase, were associated with the entire limiting membrane or subdomains within it (Figure 4C, boxes 2 and 3). Furthermore, sporadic ICD labelling that co-localised with the ECD was observed in small intra-compartmental vesicular structures, and at the periphery of the DCG itself, where strings of ILVs are normally found.

**Figure 4.**
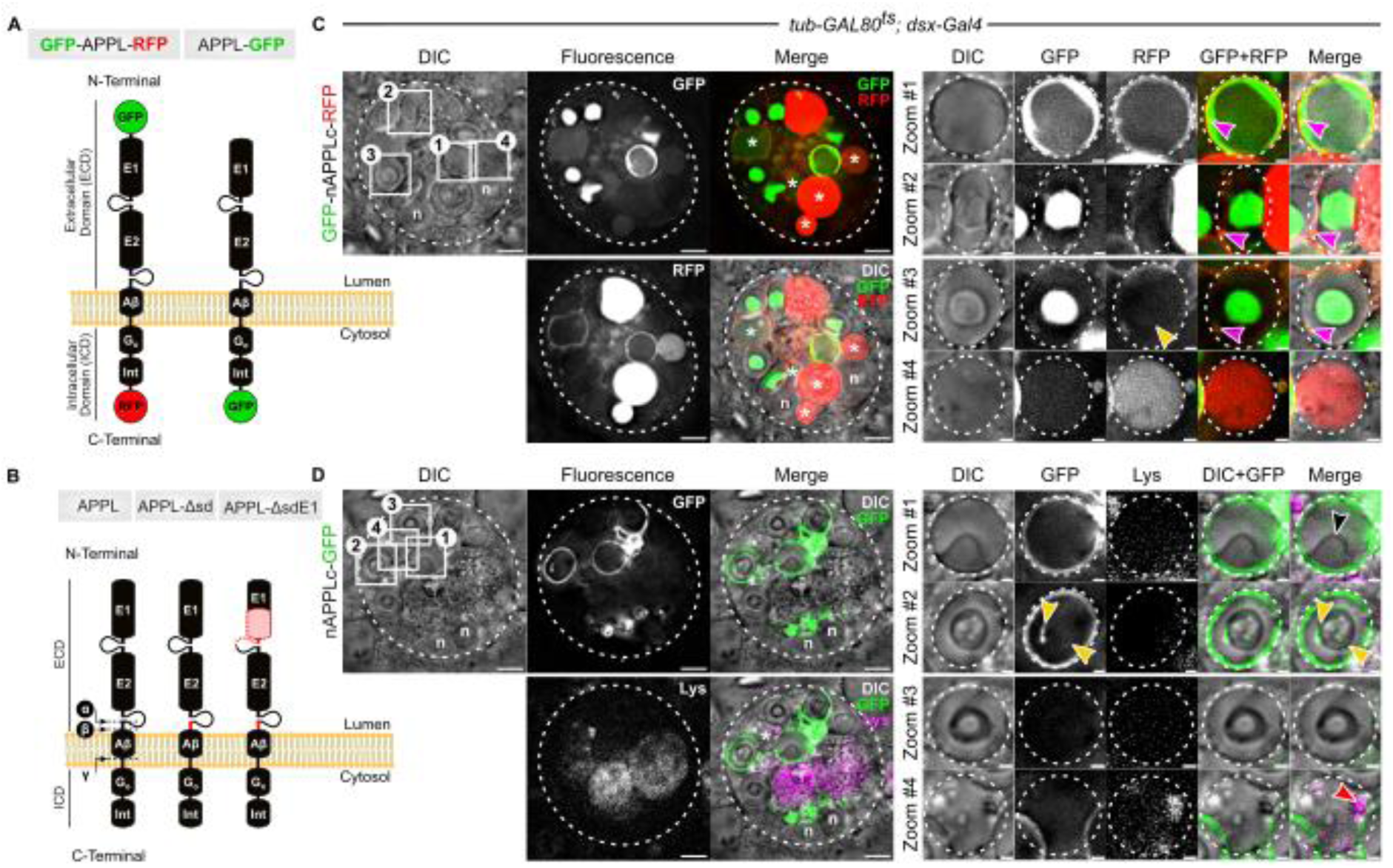
Cleavage of *Drosophila* APPL accompanies normal DCG formation. (A) Schematic showing two different fluorescently tagged APPL constructs employed in this study. (B) Schematic showing two APPL constructs employed in this study that lack α- and β-secretase cleavage sites (deletion marked in red near transmembrane domain), so the ECD cannot be released from the limiting membrane of DCG compartments or ILVs within them. The APPL-ΔsdE1 construct lacks a large proportion of the E1 extracellular domain. (C and D) *Ex vivo*, wide-field fluorescence micrographs and DIC images of SCs from 6-day-old males expressing either double-tagged APPL (dt-APPL; C) or APPL-EGFP (D). Note magnified images shown in Zoom, highlighting a DCG precursor compartment (box 1), a mature DCG compartment with APPL’s ECD (green) highly concentrated in the DCG and APPL’s ICD (red) bound to the limiting membrane (box 2), a more mature DCG compartment with the ICD no longer at its periphery (box 3) and a compartment that appears to have the DCG acidification phenotype (box 4). Other compartments with the DCG acidification phenotype are marked by a white asterisk. Yellow arrowheads mark labelled ILVs inside the DCG compartment and magenta arrowheads mark labelled ILVs at the DCG periphery. For the APPL-EGFP-labelled cell, black arrowhead (Merge) marks a DCG that has already started to form within the DCG precursor compartment and red arrowhead marks acidic microdomain. In all images, approximate cell boundary and compartment boundaries are marked with dashed white line; n = nuclei of binucleate cells; LysoTracker Red (magenta) marks acidic compartments in D. Scale bars = 5 µm and 1 µm in Zoom.

In some cells, a few compartments labelled by the ICD at their periphery contained diffuse GFP, as would be expected for compartments at early stages of the DCG acidification process (Figure 4C, box 4). There were also compartments with high levels of internalised mRFP-labelled ICD, which by DIC, had the typical ‘granular’ morphology of lysosomes (Figure 4C). This high-level accumulation may be partly explained by the observation that endosomal red fluorescent protein fusions are often disproportionately trafficked to lysosomes of SCs (eg. CD63-mCherry in Fan et al., 2020).

To confirm that the distribution of APPL’s ICD in secretory compartments was not grossly altered by the addition of a red fluorescent tag, we assessed the localisation of an APPL-EGFP C-terminal fusion in SCs (Figure 4A; Penserga et al., 2019). The EGFP-labelled ICD of APPL was observed at the limiting membrane of DCG compartments and their precursors (Figure 4D, boxes 2 and 1 respectively), mirroring the distribution of the mRFP-tagged ICD of double-tagged APPL; it was also more frequently and clearly detected in ILVs and at the periphery of DCGs. Similar to double-tagged APPL, it appeared that some, presumably more mature, DCG compartments lost the ICD from their limiting membrane (Figure 4D, box 3). There were also several compartments with the DCG acidification phenotype, where EGFP fluorescence, which is quenched in acidic compartments, was not detectable (Figure 4D, box 4).

These findings are consistent with a model in which full-length APPL traffics into Rab6-labelled DCG precursor compartments. It is then cleaved during the rapid transition period when these compartments start to generate ILVs, mini-cores and DCGs. Almost all the APPL ECD generated, and any ECD that may be subsequently trafficked into DCG compartments as they mature, becomes associated with DCG protein aggregates, which separate from the limiting membrane. The APPL ICD either remains membrane-bound or is trafficked to lysosomes. Since APPL is required for normal DCG biogenesis and its cleavage accompanies this process, we hypothesised that APPL cleavage might be a critical step in driving DCG assembly.

### *Drosophila* APPL and APPL cleavage regulate DCG protein aggregation and dissociation of these aggregates from membranes

α-, β- and γ-secretases are the key enzymes that cleave APP, separating the ECD from the ICD (Figure 4B). We therefore knocked down each of these proteases in adult SCs to assess whether they also affected DCG compartment maturation (Figure 5A). None of the knockdowns altered DCG compartment number (Figure 5B). However, DCG morphology was affected by knockdown of *β-secretase* and to a lesser extent, *γ-secretase*; some DCGs were deformed in structure, while for *β-secretase* knockdown, other DCGs lacked GFP in their centre (Figures 5A, 5C and S4A). Knockdown of each of the three secretases increased the number of large compartments with the DCG acidification phenotype (Figure 5D), suggesting that the DCG compartment maturation process is aborted more frequently in these genetic backgrounds, targeting the compartments for lysosomal degradation. Overall, this suggests that the appropriate balance of α-, β- and γ-secretase activities is critical in regulating the proper maturation of DCG compartments in SCs. However, because these proteases have targets other than APPL, the resulting knockdown phenotypes may not be explained by aberrant APPL cleavage.

**Figure 5.**
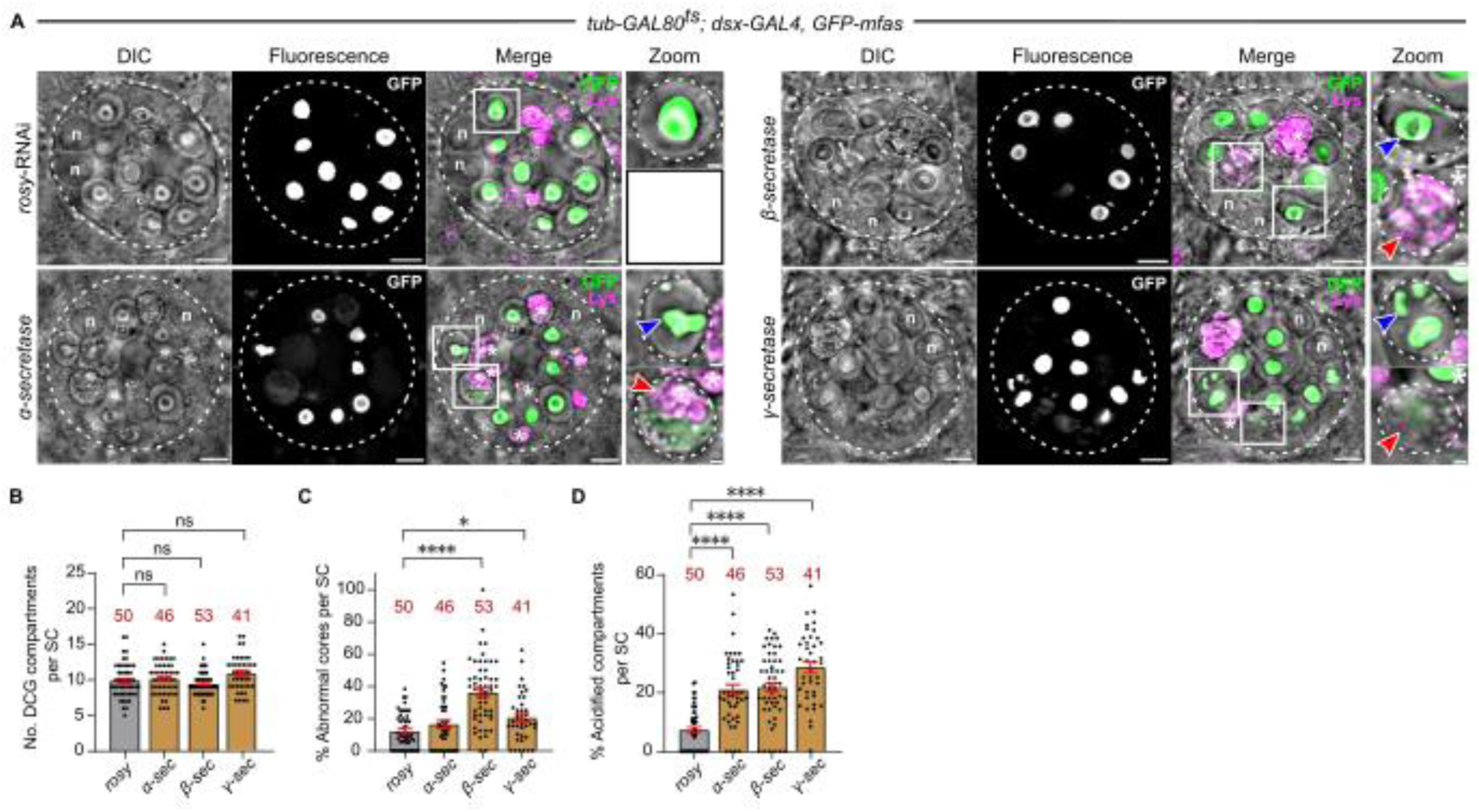
Secretases involved in *Drosophila* APPL processing regulate DCG maturation. (A) *Ex vivo*, wide-field fluorescence micrographs and DIC images of SCs from 6-day-old males expressing *GFP-mfas* gene trap and SC-specific *rosy*-RNAi or RNAi targeting *α-*, *β-* and *γ-secretases*. Note β-secretase knockdown affects DCG morphology, so that GFP-MFAS is frequently absent from a large central region within the DCG, while other secretase knockdowns rarely affect DCG morphology (shown for compartments outlined with white boxes in upper Zoom panels). However, knockdown of all secretases increases the proportion of acidified compartments in SCs. White asterisks and white boxes marked with asterisks in the Merge channel indicate DCG compartments with acidification phenotype (lower zoom panel; red arrowheads mark acidic microdomains). (B-D) Bar charts showing that knockdown of *α-*, *β-* and *γ-secretases* does not affect the number of DCG compartments (B), but for *β-secretase* knockdown, many of the DCGs are abnormal, predominantly lacking GFP-MFAS in a large central region (C). A greater proportion of large compartments display the DCG acidification phenotype following secretase knockdown than in controls (D). In all images, approximate cell boundary and compartment boundaries are marked with dashed white line; n = nuclei of binucleate cells; LysoTracker Red (magenta) marks acidic compartments. Scale bars = 5 µm and 1 µm in Zoom. For bar charts, data are represented as mean ± SEM and were analysed using the Kruskal-Wallis test, followed by Dunn’s multiple comparisons post hoc test; n = number above bar, *P<0.05, ****P<0.0001, ns = not significant.

To further investigate the role of APPL cleavage in DCG biogenesis, we overexpressed a form of APPL that lacks the α- and β-cleavage sites (APPL-Δsd; Figure 4B; Torroja et al., 1999). As we had found for human APP-YFP and fluorescently tagged forms of APPL (Figures 3 and 4), when compared to controls, overexpression of a wild type form of APPL (APPL-WT) produced some deformed DCGs and induced significantly more compartments with the DCG acidification phenotype (Figure 6 and S4C). There was a slight reduction in the number of DCG compartments, perhaps because of this lysosomal targeting (Figure 6B).

**Figure 6.**
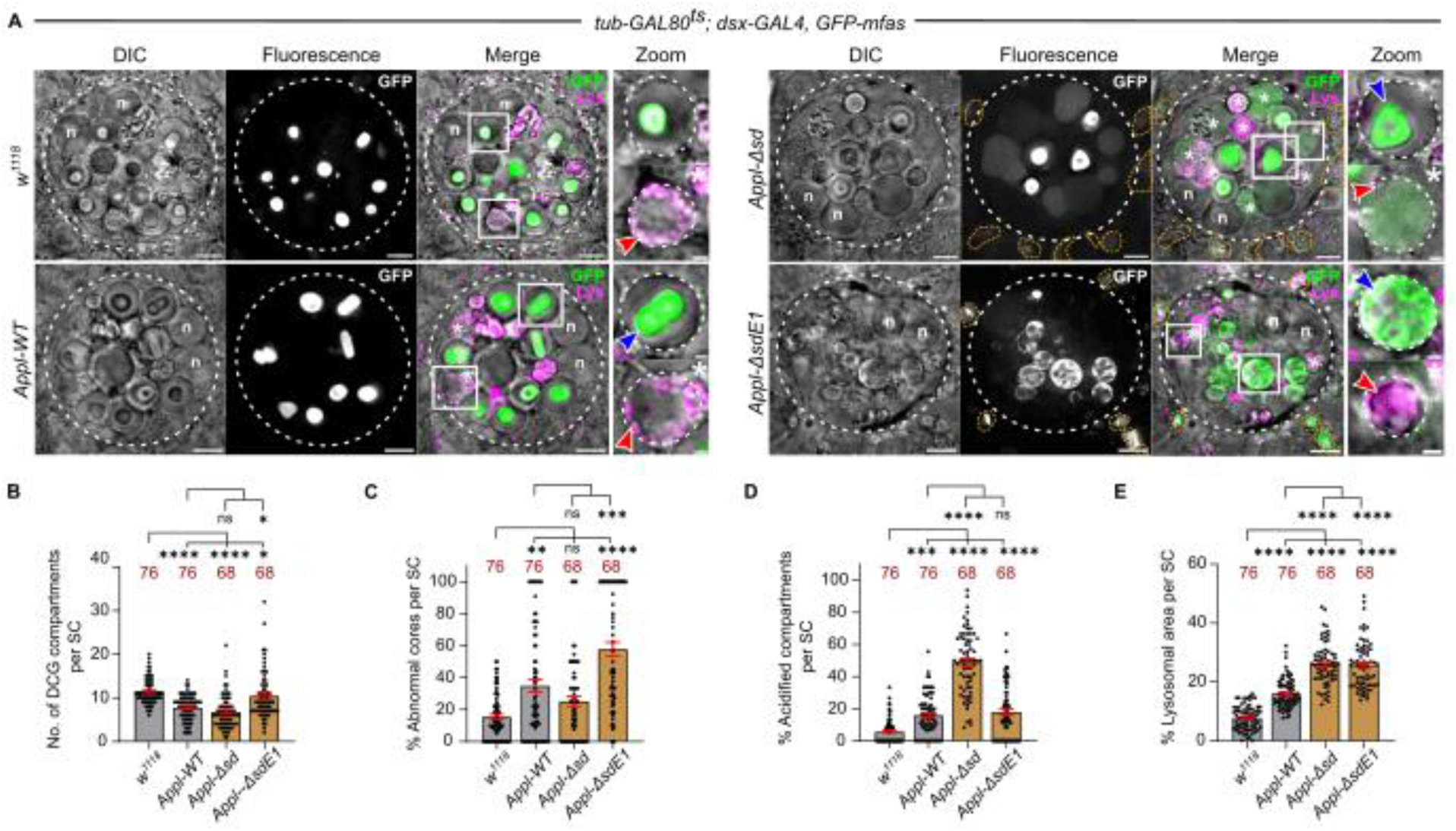
*Drosophila* APPL regulates DCG protein aggregation and its cleavage is required for normal DCG formation. (A) *Ex vivo*, wide-field fluorescence micrographs and DIC images of SCs from 6-day-old males expressing *GFP-mfas* gene trap and either no other transgene, or SC-specific wild-type APPL (APPL-WT), APPL-Δsd or APPL-ΔsdE1 (Figure 4B). Note abnormal DCG phenotypes are observed when any of the transgenes are expressed (shown for compartments outlined with white boxes in upper Zoom panels; blue arrowheads mark abnormal DCGs), but APPL-ΔsdE1 induces a unique network of primarily peripheral GFP-MFAS aggregates. White asterisks and white boxes marked with asterisks in the Merge channel indicate DCG compartments with acidification phenotype (lower zoom panel; red arrowheads mark acidic microdomains). Enlarged acidic main cell compartments containing GFP are outlined by orange dashed lines. (B-E) Bar charts showing that overexpression of APPL-WT, APPL-ΔsdE1 and particularly APPL-Δsd reduce DCG compartment number compared to controls (B). They also disrupt DCG morphology with APPL-ΔsdE1 producing the most penetrant effect (C). A greater proportion of large compartments display the DCG acidification phenotype than in *w^1118^* controls, especially for APPL-Δsd (D). Both APPL-Δsd and APPL-ΔsdE1 induce a strong and significant increase in lysosomal area (E). In all images, approximate cell boundary and compartment boundaries are marked with dashed white line; n = nuclei of binucleate cells; LysoTracker Red (magenta) marks acidic compartments in A. Scale bars = 5 µm and 1 µm in Zoom. For bar charts, data are represented as mean ± SEM and were analysed using the Kruskal-Wallis test, followed by Dunn’s multiple comparisons post hoc test; n = number above bar, *P<0.05, **P<0.01, ***P<0.001, ****P<0.0001, ns = not significant. See also Figure S4.

Overexpression of the non-cleavable APPL-Δsd mutant reduced the number of DCG compartments further, generated many more acidified DCG compartments and expanded the lysosomal area within SCs, when compared to APPL-WT overexpression (Figures 6 and S4C). Therefore, if failure to cleave APPL targets DCG compartments for lysosomal degradation. Although the APPL-Δsd protein affected DCG condensation and compartment maturation, a large central core often still formed, suggesting that DCG aggregates can dissociate from non-cleaved APPL.

In human APP, the E1 domain within the ECD (Figure 4A) is thought to play a key role in the formation of APP dimers, a process that is regulated by the divalent cations, Zn^2+^ and Cu^2+^ (August et al., 2019). Both of these ions have been implicated in DCG biogenesis (Germanos et al., 2021; Jayawardena et al., 2019). We overexpressed a non-cleavable form of APPL that lacked much of the E1 domain (APPL-ΔsdE1; Figure 4B; Torroja et al., 1999) to test the latter’s function. Remarkably, the majority of DCG compartments in cells expressing this mutant APPL protein were abnormal; in most of these defective compartments, GFP-MFAS condensed in a peripheral network, although sometimes an abnormal central DCG was also formed (Figures 6A-C, S4B and S4C). Therefore, the E1 domain of APPL appears to control the association between the APPL ECD and protein aggregates during DCG compartment maturation. In its absence, more stable membrane-associated aggregates form, preventing central DCG formation when APPL cannot be cleaved. The defective compartment maturation induced did not affect the proportion of acidified DCG compartments formed in SCs, when compared to APPL-WT-expressing cells, but the lysosomal area was significantly increased (Figures 6A, 6D and 6E), suggesting preferential targeting to lysosomes.

We also observed an additional novel phenotype in accessory glands that were overexpressing the two non-cleavable APPL proteins. In these genotypes, the main cells, the other much more numerous epithelial cell type in the accessory gland, contained high levels of internalised GFP-MFAS, which appeared to be located within unusually enlarged acidic compartments (Figures 6A, S4D and S4E). This suggests that GFP-MFAS continues to be secreted in the presence of APPL-Δsd proteins, and is either preferentially endocytosed from the accessory gland lumen by non-expressing cells and/or is not degraded normally when internalised by these cells.

Taken together, our data suggest that the absence of APPL or the failure to cleave this molecule leads to defective DCG biogenesis and in all cases, an increased proportion of these compartments is targeted for lysosomal degradation. This produces either an excess of compartments with the DCG acidification phenotype or an expansion of lysosomal compartments. When APPL is not cleaved, this also affects the uptake and/or breakdown of secreted MFAS by other cells.

### Amyloidogenic Aβ-42 peptides disrupt DCG biogenesis in SCs

Having identified novel functions for APPL and its cleavage in DCG protein aggregation, aggregate release from membranes and DCG compartment maturation and trafficking, we hypothesised that these processes might be selectively misregulated in Alzheimer’s Disease. To test this in our model, we overexpressed in SCs a wild-type form of the pathological Aβ-42 peptide that is generated from APP following cleavage by β- and γ-secretases (O’Brien et al., 2011), and two mutant forms of this peptide with elevated amyloidogenic activity that lead to familial Aβ-peptide-associated amyloid disease, the Dutch (Van Broeckhoven et al., 1990) and Iowa (Grabowski et al., 2001) mutants (Van Nostrand et al., 2002) (Figure 7B). In *Drosophila*, expression of these molecules in either the nervous system or the eye induces neurodegenerative phenotypes in adults, with wild type Aβ-42 producing the mildest phenotypes (Chouhan et al., 2016; Metsla et al., 2022).

**Figure 7.**
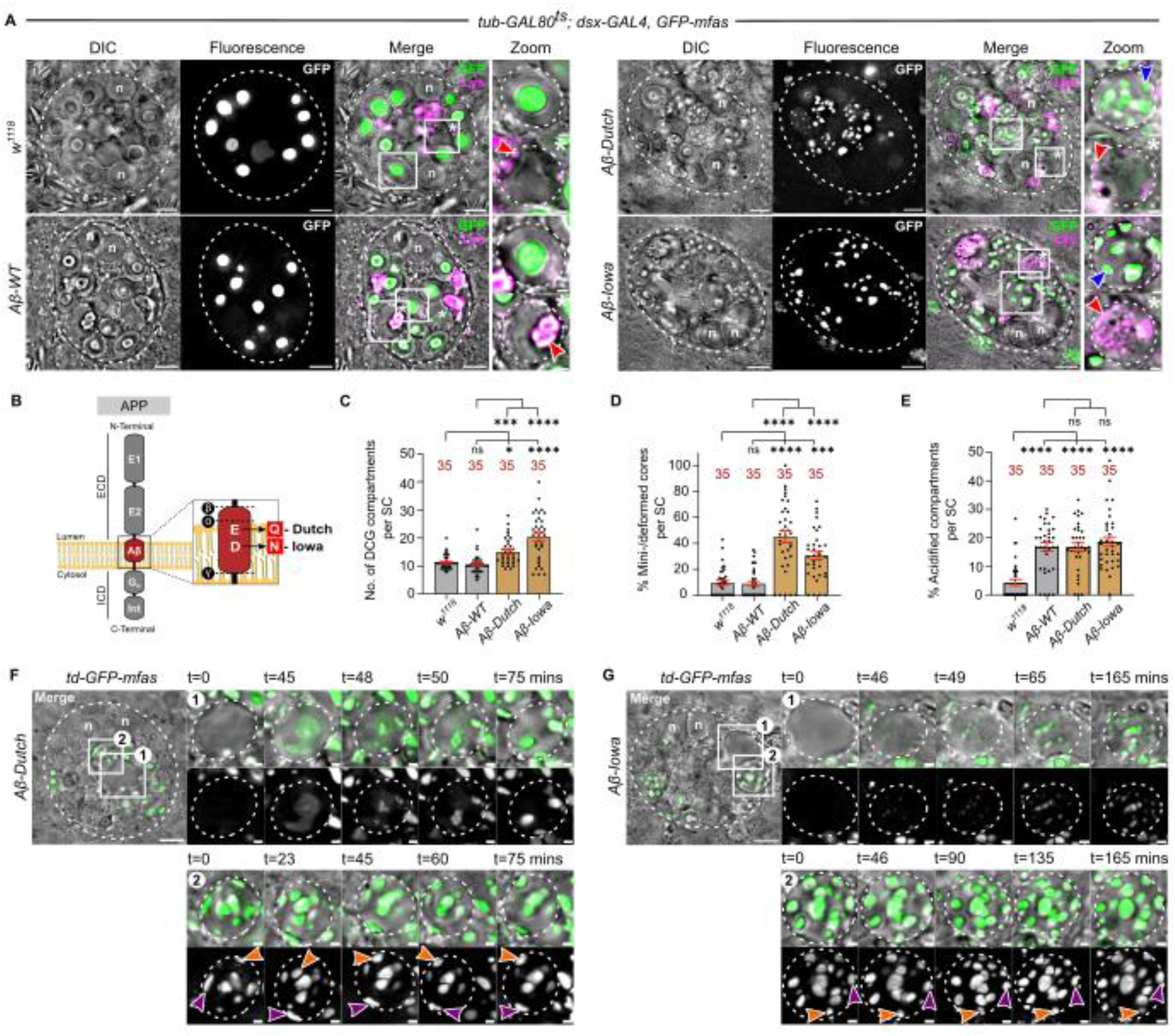
Expression of pathological mutant Aβ-42 peptides in SCs disrupts DCG biogenesis and increases lysosomal targeting of DCG compartments. (A) *Ex vivo*, wide-field fluorescence micrographs and DIC images of SCs from 6-day-old males expressing *GFP-mfas* gene trap and either no other transgene, or wild type Aβ-42 peptide, or either the Iowa or Dutch mutant Aβ-42 peptides. Note mini-core phenotypes with Iowa and Dutch mutants, and DCG acidification phenotype induced by overexpression of all Aβ-peptides. White asterisks and white boxes marked with asterisks in the Merge channel indicate DCG compartments with acidification phenotype (lower zoom panel; red arrowheads mark acidic microdomains). (B) Schematic showing different Aβ-42 peptides expressed in this study. (C-E) Bar charts showing that overexpression of Aβ-42 Dutch and Aβ-42 Iowa mutant peptides increases DCG compartment number compared to controls (C) and generates a mini-core phenotype in many of these compartments (D). All Aβ-peptides induce a greater proportion of large compartments displaying the DCG acidification phenotype than in *w^1118^* controls (E). (F, G) Stills from time-lapse movies of DCG biogenesis in SCs from 6-day-old males expressing *GFP-mfas* gene trap and Dutch (F) or Iowa (G) Aβ-42 mutant, focusing on one compartment as it forms mini-cores (white box 1) and another mature compartment (white box 2). Note that multiple GFP-MFAS-containing mini-cores form, but they are relatively immobile (blue arrowheads) and unlike control cells (Figure 1H), they do not fuse. The DCG compartments are also less motile within the cell compared to controls. For the Aβ-42 Iowa mutant, condensation appears to take place extremely rapidly and no GFP-MFAS clouds are observed prior to this event. In all images, approximate cell boundary and compartment boundaries are marked with dashed white line; n = nuclei of binucleate cells; LysoTracker Red (magenta) marks acidic compartments in A. Scale bars = 5 µm and 1 µm in Zoom. For bar charts, data are represented as mean ± SEM and were analysed using the Kruskal-Wallis test, followed by Dunn’s multiple comparisons post hoc test; n = number above bar, *P<0.05, ***P<0.001, ****P<0.0001, ns = not significant. See also Figure S5, and Movies S5 and S6.

Overexpression of all of these molecules in SCs disrupted aspects of DCG formation (Figures 7A and 7C-E). While wild type Aβ-42 did not affect the total number of DCG compartments and intact DCGs per cell, more DCG compartments were formed in SCs expressing the Dutch and Iowa Aβ-42 mutants and more than one third of these contained mini-cores, often in addition to a central abnormally shaped small DCG (Figures 7A, 7C and 7D). An increased number of acidified DCG compartments was also observed, even with wild type Aβ-42 overexpression (Figures 7A and 7E). Time-lapse movies of DCG compartment maturation in SCs expressing either the Dutch or Iowa Aβ-42 mutant peptides revealed a very similar phenotype to *Appl* knockdown with static peripheral mini-cores within compartments that also appeared to have restricted motility within the cell (Figure 7F and 7G; Movies S5 and S6).

The intracellular phenotypes observed in SCs emerge after only 6 days of transgene overexpression. During this time, many mature DCG compartments normally fuse with the secondary cell’s apical membrane and release their contents into the lumen of the accessory gland (Redhai et al., 2016). The lumen expands rapidly during the first three days post-eclosion, a process primarily driven by secretion from the much more abundant main cells. We assessed the lumenal accumulation of GFP-MFAS over six days post-eclosion and checked whether expression of Aβ-42 in SCs led to any obvious effects on GFP levels or extracellular GFP-MFAS aggregation. In wild type six-day-old males, relatively homogeneously distributed GFP-MFAS was observed at high levels in the lumen of the AG (Figure S5A), consistent with our previous observations that DCGs are rapidly dispersed following secretion (Redhai et al., 2016). The lumenal distribution of GFP-MFAS in males where SCs were expressing wild type and mutant Aβ-42 was not obviously affected and indeed, the total levels of secreted GFP-MFAS appeared to be increased by Aβ-42 expression (Figure S5). We therefore conclude that in the fly model, mutant Aβ-peptides modulate intracellular DCG protein aggregation events and overload the endolysosomal system prior to any notable accumulation of abnormal extracellular aggregates.

## Discussion

APP cleavage products are thought to be central players in protein aggregation events leading to amyloid formation in AD. However, APP’s roles in the early stages of this disease remain unclear. Changes in secretion and endolysosomal trafficking are well-established early features of AD, which can modulate APP cleavage (Kimura and Yanagisawa, 2018). Conversely, loss of APP function also affects processes associated with secretion (Nalivaeva and Turner, 2013). However, determining how these changes might initiate pathological defects at the subcellular level is challenging, because it requires the analysis of events taking place inside compartments that are typically at the limit of light microscope resolution.

We have overcome this problem by studying APP function *ex vivo* in a *Drosophila* cell type that has highly enlarged secretory and endosomal compartments. We demonstrate that APPL, the fly APP homologue, has a highly specific physiological function in regulating protein aggregation events required to form a large DCG. Normal DCG biogenesis requires transmembrane APPL, but the process is also dependent on release of the APPL extracellular domain by proteolytic cleavage. If this latter event fails, APPL expression is suppressed, or pathological Aβ-42 peptides are expressed in SCs, DCG biogenesis and compartment maturation are disrupted. This leads to defective compartment trafficking and targeting for lysosomal degradation, mirroring early AD-associated phenotypes commonly observed in neurons (Kimura and Yanagisawa, 2018).

In studying DCG biogenesis, we also discovered a role for the TGFBI homologue, MFAS, which is essential for all aspects of SC DCG protein aggregation. Mutant forms of this fibrillar protein lead to the human amyloidogenic disease, corneal dystrophy. We have demonstrated a co-dependence between APPL and MFAS in physiological protein aggregation, with APPL acting as a membrane-associated regulator of MFAS condensation and amyloidogenic forms of APP modulating this event. Cross-seeding between amyloid proteins has been observed experimentally (Subedi et al., 2022). It has been proposed as an explanation for higher than expected co-morbidities of amyloidogenic diseases (Spires-Jones et al., 2017). Our findings suggest that at least in some cases, there may be a physiological basis for such effects.

### MFAS, the TGFBI homologue, drives protein aggregation in SC DCGs

Using real-time imaging of the *GFP-mfas* gene trap and the large DCG precursor compartments in SCs, we visualised the protein condensation events that lead to formation of a very large (approximately 3 μm diameter) DCG. Often, a cloud of GFP-MFAS protein rapidly aggregates into a central core via a non-homogeneous intermediate. However, in other cases, many mini-cores initially form at the periphery of the compartment, which rapidly become motile and fuse together.

Like control cells, *GAPDH2* knockdown SCs, which were previously shown to contain defective DCG compartments with many peripheral mini-cores surrounded by ILVs (Dar et al., 2021), assemble multiple mini-cores near the compartment’s limiting membrane. These mini-cores and the DCG compartments are motile, but the mini-cores only rarely collide, and when they do, they do not appear to be able to fuse. GAPDH2 is required for normal clustering of ILVs in SCs, forming ILV chains that extend from the limiting membrane to the central DCG. One explanation for our findings is that in the absence of these radial ILV chains, ILV-associated mini-cores are unable to come together to form a central DCG.

Knockdown experiments revealed that MFAS, an extracellular matrix protein that was originally reported to be membrane-associated (Hu et al., 1998), is essential for both mini-core and large DCG biogenesis (Figure 1). The human MFAS homologue, TGFBI, is structurally very similar to MFAS, except that it includes a C-terminal RGD domains that provide extra integrin-binding motifs (Nielsen et al., 2020). Mutations in *TGFBI* are responsible for most cases of autosomal dominant corneal dystrophy, where the protein forms extracellular amyloid deposits (Han et al., 2016). Interestingly, human APP and TGFBI are co-expressed in the secretory pathway of corneal cells (Choi et al., 2019) and associate in putative amyloid deposits associated with calcific aortic valve disease (Heuschkel et al., 2020).

Our work suggests that TGFBI’s *Drosophila* homologue has a natural propensity to rapidly form large super-molecular aggregates under specific microenvironmental conditions. These conditions are met when large Rab6-positive DCG precursor compartments formed at the *trans*-face of the Golgi receive input from Rab11-positive recycling endosomes (Wells et al., 2023). It seems likely that part of the condensation process involves changes in pH and/or ion concentrations, which have previously been implicated in DCG biogenesis (Wu et al., 2001; Li, 2014; Yoo and Albanesi, 1990). However, our real-time imaging studies indicate that the concentration of GFP-MFAS may also increase during the hour before DCGs are formed (see Movies S1 and S2), suggesting that the trafficking of MFAS to precursor compartments may also be critical in driving protein aggregation.

It will now be interesting to assess whether TGFBI proteins carrying mutations associated with corneal dystrophy, some of which are in residues conserved in MFAS, promote or interfere with DCG biogenesis. One other parallel that also requires further investigation relates to new multimolecular extracellular signalling nanoparticles called supermeres, which contain a diverse range of proteins and RNAs (Zhang et al., 2021). TGFBI is the most abundant protein in these structures, which also include the cleaved extracellular domains of proteins like APP, as well as extracellular proteases and GPI-anchored proteins. Here, we have demonstrated the presence of APP’s ECD in SC DCGs (Figure 4C). The extracellular protease <u>an</u>giotensin I-<u>c</u>onverting <u>e</u>nzyme (ANCE) also accumulates in SC DCGs (Rylett et al., 2007; Corrigan et al., 2014) and we have previously used a GPI-anchored form of GFP as a specific DCG marker (Redhai et al., 2016), suggesting that SC DCGs share structural similarities to supermeres. Supermeres might therefore form in maturing secretory compartments through interactions with endosomal and exosomal membranes, as we have found for DCGs. Furthermore, since DCGs provide the storage site for signalling molecules, it could explain why supermeres have elevated signalling activity (Zhang et al., 2021).

### APP regulates protein aggregation events and its cleavage is required for normal DCG assembly and secretory trafficking

Rab11-exosomes are a novel subtype of exosomes formed in compartments labelled by the recycling endosomal marker Rab11 (Fan et al., 2020), rather than by late endosomes, which were previously thought to be the key source of these secreted vesicles. Even though Rab11-exosomes may make up only a small proportion of cellular EV output, when their biogenesis is selectively inhibited, specific defects in physiological and pathological cell communication are observed (Marie et al., 2023). In SCs, Rab11-exosomes form in regulated secretory compartments, where they have an additional role, as evidenced by *GAPDH2* knockdown, co-ordinately controlling with endosomal membranes the normal protein aggregation events required for large DCG biogenesis.

Our analysis of APPL function has provided an explanation for the link between membranes and protein aggregation in SCs (Marie et al., 2023; Wells et al., 2023; Figure 8). APPL is the only APP-like homologue in flies; it is expressed at high levels in the nervous system, but is also transcribed in other tissues (Leader et al., 2018). Analysis of APPL trafficking and cleavage in SCs with a double-tagged protein revealed that intact APPL localises to the limiting membrane of Golgi-derived, DCG precursor compartments marked by Rab6, but as soon as the Rab6 to Rab11 transition occurs, and ILVs and DCGs begin to form, cleavage occurs. APPL’s ECD associates with the DCG, and the membrane-linked ICD either remains on the limiting membrane of DCG compartments and on ILVs or is targeted to lysosomes (Figure 4C). Overexpressing APPL or APP directs more DCG compartments towards lysosomes, with particularly severe phenotypes observed when the ECD cannot be released from membranes, leading to a build-up of large lysosome-targeted compartments (Figure 6).

**Figure 8.**
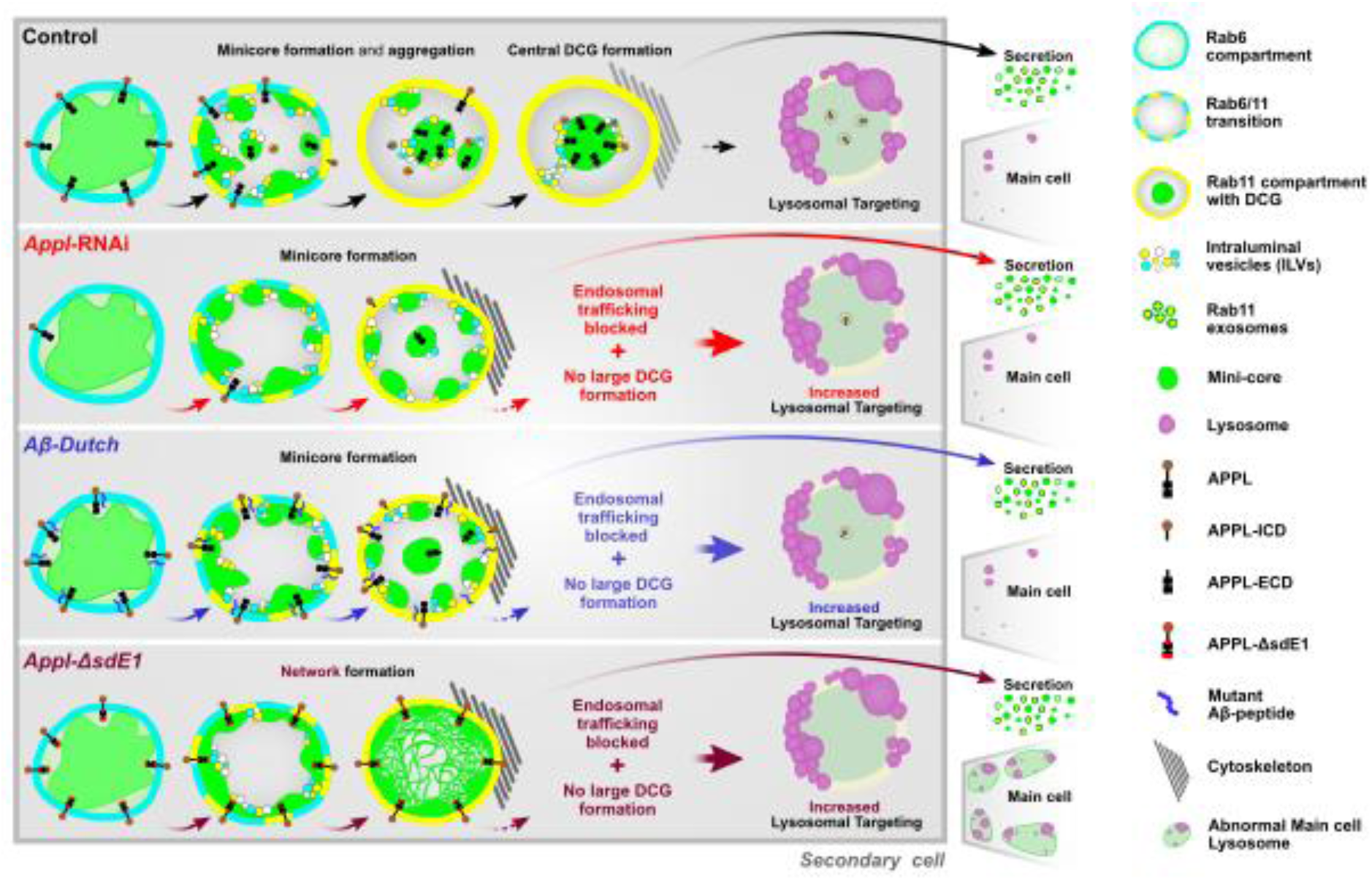
Schematic model of APPL-regulated DCG formation in *Drosophila* secondary cells. Schematic shows DCG assembly process in control conditions (top row) with large central DCG (green) forming from peripheral mini-cores that interact with both the compartment’s limiting membrane and ILVs. This initial step is triggered by a Rab6 to Rab11 transition. In some compartments, it is not possible to distinguish the biogenesis of individual mini-cores, although the central DCG still forms by condensation of a cloud of diffuse MFAS, which is initially in contact with the compartment’s limiting membrane. *Appl* knockdown or pathological (Dutch) mutant Aβ-42 expression suppresses coalescence of mini-cores, increases lysosomal targeting, but also suppresses secretory compartment motility within the secondary cell, presumably by stabilising cytoskeletal interactions. Non-cleavable forms of APPL alter the properties of secretions, so that MFAS is more readily endocytosed by main cells and leads to lysosomal enlargement in these cells. The APPL--ΔsdE1 mutant also promotes limiting membrane-associated MFAS condensation, producing an aggregated network at the compartment’s periphery.

Reducing or removing *Appl* does not block all DCG protein aggregation, but aggregates are frequently associated with the limiting membrane of DCG compartments, either as mini-cores or as a single large core (Figures 3 and S3). These compartments are also preferentially targeted for lysosomal degradation, indicating that APPL is critical for normal DCG maturation in SCs. Furthermore, non-acidic DCG compartments that contain mini-cores are visibly less motile within the cytoplasm than normal compartments, perhaps because of more stable links with the cytoskeleton. Finally, the mini-core phenotype induced by *Appl* knockdown is strongly suppressed by human APP, suggesting that the aggregation function of APPL is evolutionarily conserved.

What mechanisms might target DCG compartments for lysosomal degradation, either when APPL levels are reduced or elevated, or when APPL fails to be cleaved? In many examples, such as *Appl* knockdown or overexpression of non-cleavable APPL, DCG aggregates remain abnormally associated with the limiting membrane of compartments. When non-cleavable APPL mutant proteins are overexpressed, a proportion of the many acidified DCG compartments formed can fuse to lysosomes, increasing the lysosomal area. However, the mini-core phenotype observed following *Appl* knockdown has highlighted that defective DCG compartments, and the membrane-associated mini-cores within them, have reduced motility, which presumably suppresses compartmental trafficking to large lysosomes in the cell. Therefore, a failure to separate protein aggregates from the DCG compartment’s limiting membrane may have the dual effect of disrupting compartment maturation while preventing these compartments from being trafficked normally for degradation.

This immobilisation model may not explain the high levels of the DCG acidification phenotype in all genotypes. Most notably, α-, β- and γ-secretase knockdowns do not appear to affect the association of GFP-MFAS aggregates with the compartment’s limiting membrane, but are associated with elevated DCG compartment acidification (Figure 5). However, these secretases are known to affect the processing of other proteins in the secretory and endosomal system, so it remains unclear whether the phenotypes observed result from reduced APPL cleavage.

APPL not only controls the maturation of DCG compartments and dissociation of membrane:aggregate complexes, it also regulates MFAS-mediated protein aggregation itself. Overexpressing non-cleavable full-length APPL does not induce defective protein aggregation along the entire limiting membrane of DCG compartments (Figure 6A), where this molecule will be located. This suggests that if membrane-associated ECD has priming activity for aggregation, it does not irreversibly attach the ECD to those aggregates. However, the unique peripherally concentrated network of GFP-MFAS aggregation induced by non-cleavable APPL lacking a functional extracellular E1 domain (Figure 6A) strongly suggests that the E1 domain modulates aggregate priming activity. In human APP, the E1 domain is involved in divalent cation-regulated APP dimerization (August et al., 2019), and also undergoes a conformational change at acidic versus neutral pH (Hoefgen et al., 2015). These microenvironmental changes are both involved in regulating DCG aggregation (Germanos et al., 2021; Jayawardena et al., 2019; Yoo and Albanesi, 1990; Wu et al., 2001). The E1 domain therefore provides a potential sensor for microenvironmental regulation of protein aggregate priming that could be critically relevant to pathologies that disrupt aggregation.

Based on our findings, we propose a model for APPL function in DCG biogenesis, which involves roles both in regulating protein aggregation and releasing those aggregates from membranes associated with their compartment of origin, an event requiring proteolytic separation of the APPL ECD (Figure 8).

### Pathological Aβ-peptides disrupt DCG biogenesis and compartment motility

Expression in SCs of mutant Aβ-42 peptides that are known to induce neurodegenerative pathology in mammals and flies induced defects in DCG compartment maturation (Figure 7A), which phenocopied changes in SCs with reduced APPL function. Most notably, mini-cores were associated with the periphery of DCG compartments, which were more frequently acidified, but also immobilised. Defective neuronal endolysosomal trafficking is one of the earliest phenotypes reported to be associated with AD (Kimura and Yanagisawa, 2018), and our study suggests that this may be linked to dysfunction of a physiological APP-dependent process in secretion.

We have previously observed that DCG compartments occasionally fuse with large lysosomes in normal SCs (Corrigan et al., 2014) and in our current study, we found that sporadic compartments in control cells display the DCG acidification phenotype (Figure S1G). The acidification phenotype induced by *Appl* knockdown or Aβ-peptide expression is much more common than in control cells, perhaps because the defective compartmental trafficking in these cells blocks degradation of defunct compartments. If these compartments are ultimately secreted, the contents may also be dissipated abnormally. Indeed, SCs overexpressing non-cleavable forms of APPL, which also induce increased DCG acidification, secrete GFP-MFAS that accumulates in other cells within the accessory gland and appears to affect lysosomal degradation. This suggests that the mis-aggregation of proteins and lysosomal defects induced in SCs can be transferred to cells that do not normally make MFAS or DCGs.

One phenotype that we have not observed in Aβ-peptide-expressing SCs is cell death. We have previously found that these post-mitotic cells are remarkably robust, since viable cells can completely change their morphology, delaminate from the gland epithelium and even be transferred to females upon mating (Leiblich et al., 2012). To date, we have not observed apoptotic SCs, even after 14 days of Aβ-peptide expression. It will be interesting to follow SC fate for extended periods of time in the future to determine whether the phenotypes observed become more severe and whether cells may ultimately start to die. Importantly, the cellular defects we observe also appear to emerge much more rapidly than any defects in the lumen of the accessory gland into which SCs secrete, consistent with the idea that the DCG maturation defects could be an early pathological phenotype.

### APPL’s subcellular roles in SCs and their relationship to APP functions in neurons

Studying APPL and APP functions in SCs has provided unique opportunities to dissect out the activities of these molecules at the sub-compartmental level, because SC secretory compartments are so highly enlarged. But are these functions relevant to understanding the role of APP and the effects of Aβ-peptides in neurons or indeed, any other cell type?

When APP or APPL is expressed in other *Drosophila* cell types, such as developing neurons and secretory cells of the salivary gland, it is trafficked through the regulated secretory pathway and can be processed in these secretory compartments (Ramaker et al., 2016; Neuman et al., 2021). Loss of function of APPL in *Drosophila* neurons is viable, but leads to learning and behavioural phenotypes, and defects in neuronal outgrowth and synaptogenesis (Luo et al., 1992; Mora et al., 2013; Torroja et al., 1999). Other experiments have indicated that there may be parallels between loss-of-function *Appl* phenotypes and those phenotypes induced by Aβ-peptides (Singh et al., 2017). Indeed, several lines of evidence support the idea that Aβ-peptide-induced effects in tissues and associated neurodegeneration in humans represent APP loss-of-function phenotypes linked to altered secretory activity of neurons (Gouras et al., 2013; Kepp, 2016; Kim and Bezprozvanny, 2023).

Could the Rab6/Rab11-regulated DCG pathway characterised in SCs play a role in early neuronal events associated with APP function and AD? Although many models for APP processing propose an important role for APP trafficking to the cell surface prior to cleavage, subcellular localisation of the ECD and ICD of APP and APPL suggest that cleavage can take place within secretory and endosomal compartments (Muresan et al., 2015; Ramaker et al., 2016). Rab6 is reported to play a role in regulating the proteolytic processing of APP (McConlogue et al., 1996), as is the adaptor protein complex AP-1 (Januário et al., 2022), which is essential for the trafficking that controls DCG biogenesis in SCs (Wells et al., 2023).

Furthermore, several lines of evidence indicate a role for Rab11-directed trafficking in the cleavage of APP and formation of Aβ-peptides (Udayar et al., 2013; Sultana and Novotny, 2022), and support the idea that these processing events take place in maturing synaptic vesicles (Groemer et al., 20011; Del Prete et al., 2014). There is a known role for Munc18-interacting proteins (Mints), adaptor proteins that bind to the endocytic sorting motif of APP, in the secretion of Aβ-peptides from mammalian neurons (Ho et al., 2008; Sullivan et al., 2014), while in flies, APPL/Mint interactions appear to be important in synapse formation (Ashley et al., 2005). Both Mint1 and Munc-18 have also been implicated in dense-core granule secretion in neuroendocrine cells (Schütz et al., 2005).

Another important parallel relates to the potential role of exosomes in protein aggregation and APP-mediated functions in flies and humans. In *Drosophila*, larval neuromuscular junction establishment, which is regulated by APPL (Ashley et al., 2005), is also controlled by Rab11-dependent exosomes (Koles et al., 2012), consistent with the link we have identified. Furthermore, there is increasing evidence that exosomes have a role in Aβ-peptide secretion, propagation and plaque formation (Sardar Sinha et al., 2018; Kaur et al., 2021). The endosomal compartment in which these exosomes might be formed is yet to be fully characterised (Edgar et al., 2015). Our study suggests a role for Rab11-exosomes, which is supported by the observation that Rab11-exosome markers may be enriched in the CSF secretome from AD patients (Li et al., 2022; Table S1). In fact, the link between ILVs in endosomal compartments and protein aggregation is not unique to APP, Aβ and TGFBI. Endolysosomal ILVs are required for the priming of amyloid formed by the protein PMEL in melanocytes, a process also involving Apolipoprotein E, an important player in AD susceptibility (van Niel et al., 2015).

### SCs as a model for AD pathology

Our studies in SCs suggest that cleavage of APP and dissociation of protein aggregates from compartment membranes are critical steps in the normal maturation of DCG compartments. If these steps fail, either because APPL levels are modulated or the protein is not properly processed, or by overexpressing mutant Aβ-42 peptides, DCG compartments are more frequently targeted for lysosomal degradation. In normal SCs, lysosome-targeted DCG compartments subsequently fuse with and are degraded by a large lysosome. By contrast, when APPL-dependent events fail to take place normally in SC compartments, DCG compartments can lose mobility and acidified compartments accumulate. Disruption of endolysosomal trafficking and degradation has been proposed as a mechanism for generating AD pathology (Peric and Annaert, 2015; Kimura and Yanagisawa, 2018). It will now be important to test other AD risk genes in the SC system, and to identify additional players regulating the maturation of DCG compartments in SCs, which might also be associated with AD.

We initially tested the role of APPL in DCG biogenesis, because of the overlap between the proteomic CSF signature for AD and the Rab11a-exosome proteome. This parallel suggests that there may be enhanced release of Rab11a-exosomes and their associated proteins in AD patients, even though the secretory compartments involved may be more frequently diverted towards lysosomal degradation in the genetic backgrounds we have investigated. Our findings in SCs do not preclude an increase in secretory trafficking. Indeed, despite the endolysosomal phenotype, the levels of GFP-MFAS in the accessory gland lumen appear to be increased following Aβ-42 expression. One possible explanation is that many of the defective DCG compartments are still ultimately secreted. In addition, perhaps SCs compensate for the defects in secretory trafficking by increasing traffic through the secretory pathway, a change that may also take place in neurons of AD patients.

The *Drosophila* SC model offers new opportunities to study in detail specific events in pathological amyloidogenesis in an *in vivo* system. For example, we noticed that the Iowa Aβ-42 mutant peptide seemed to induce more rapid mini-core formation compared to the other Aβ-42 peptides tested or to wild type cells, suggesting possible differences in the amyloid-inducing activity of this mutant that could be explored further. GAPDH has been suggested to act as a seed for Aβ amyloidogenesis (Gerszon and Rodacka, 2018), a proposal that can now be tested *in vivo* in flies. Finally, increasing the level of APPL-dependent, membrane:aggregate association by overexpressing non-cleavable forms of APPL in SCs induces spreading of aggregates between cells in the accessory gland. It will be interesting to study this process further to determine whether it in any way mirrors amyloid spreading events in the brain in AD.

Perhaps the most surprising finding in our study is that homologues of two human proteins involved in amyloidogenic disease, APP and TGFBI, co-operate together under physiological conditions to induce protein aggregation via reversible interaction with membranes. Cross-seeding between amyloid proteins is observed experimentally (Subedi et al., 2022) and has been proposed as an explanation for higher than expected co-morbidities of amyloidogenic diseases (Spires-Jones et al., 2017). The possibility that some of these proteins may normally control each other’s aggregation might provide clues regarding the biological basis of these phenomena and highlight a more general role for exosomes and membranes in these processes.

It has previously been noted that other neurodegenerative diseases might share with AD some cell and molecular defects that contribute to disease (Tofaris and Buckley, 2018). In this regard, Parkinson’s Disease is commonly associated with endolysosomal trafficking phenotypes, while CHMP2B and CHMP1B, whose homologues both play roles in Rab11-exosome and DCG biogenesis in flies (Marie et al., 2023), are linked to frontotemporal dementia and hereditary spastic paraplegia respectively (Goedert et al., 2012; Reid et al., 2005). It will therefore be interesting to investigate whether one of the common mechanisms inhibited in these diseases is the dissociation of protein aggregates from endosomal and exosomal membranes.

## Materials and Methods

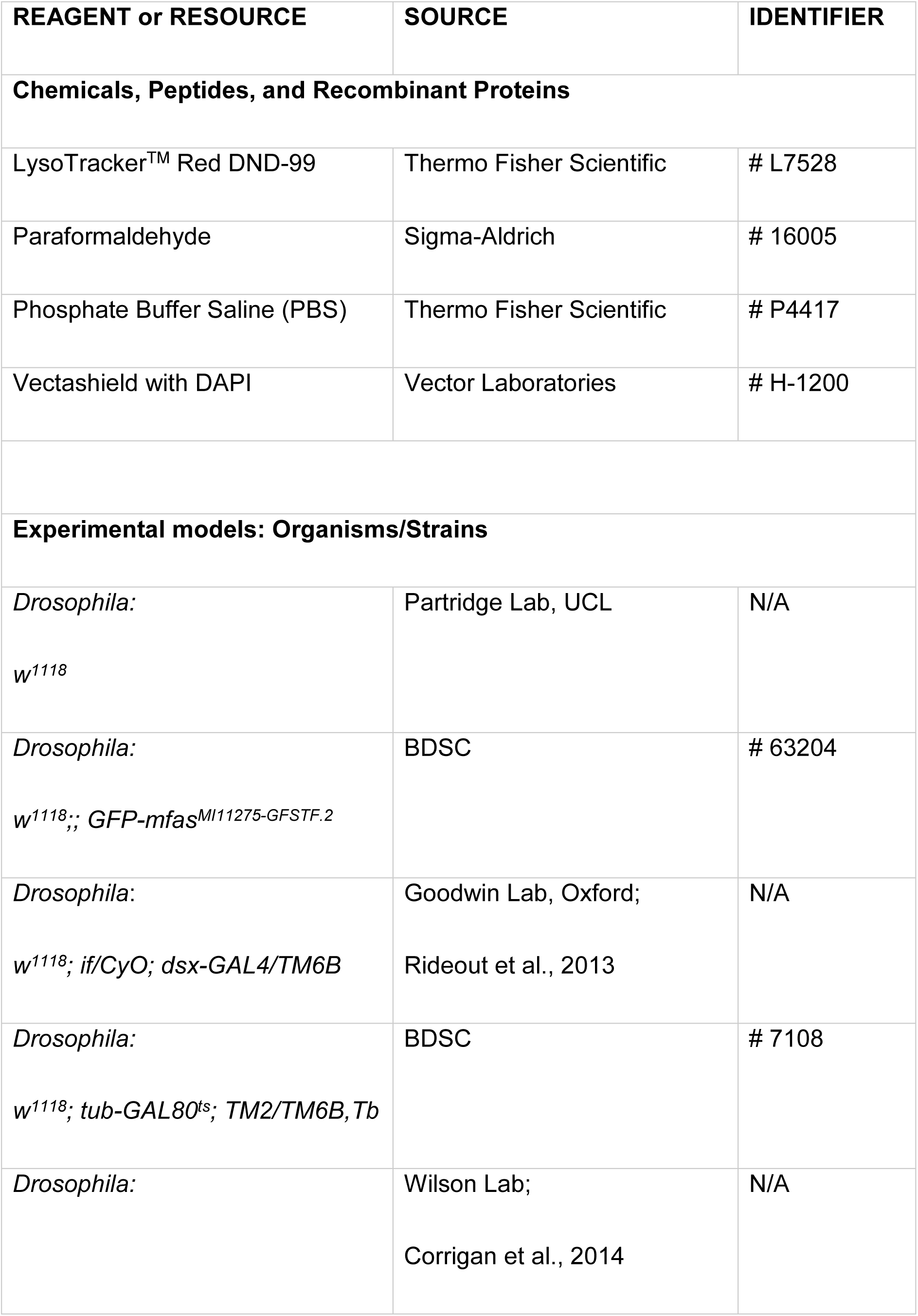
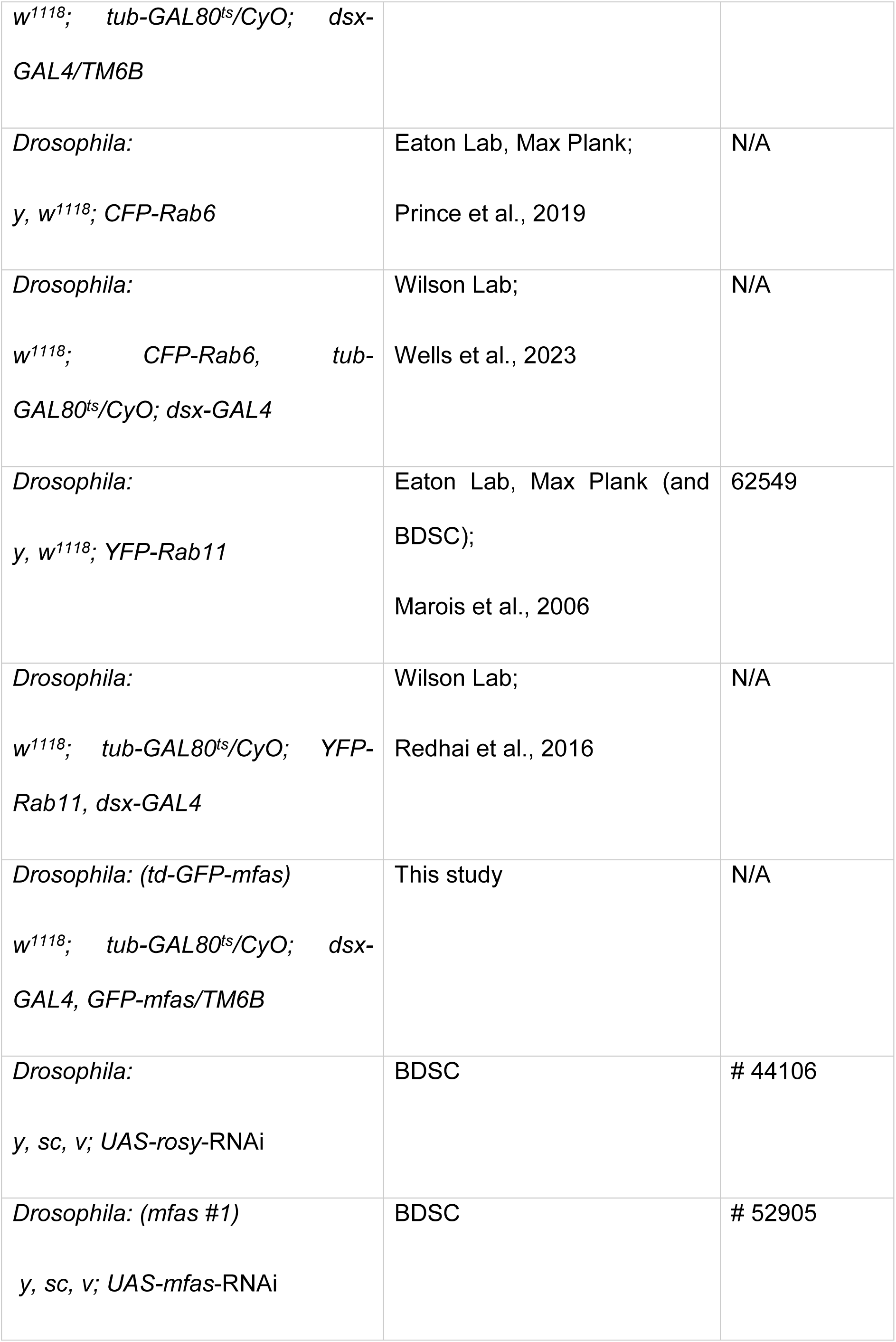
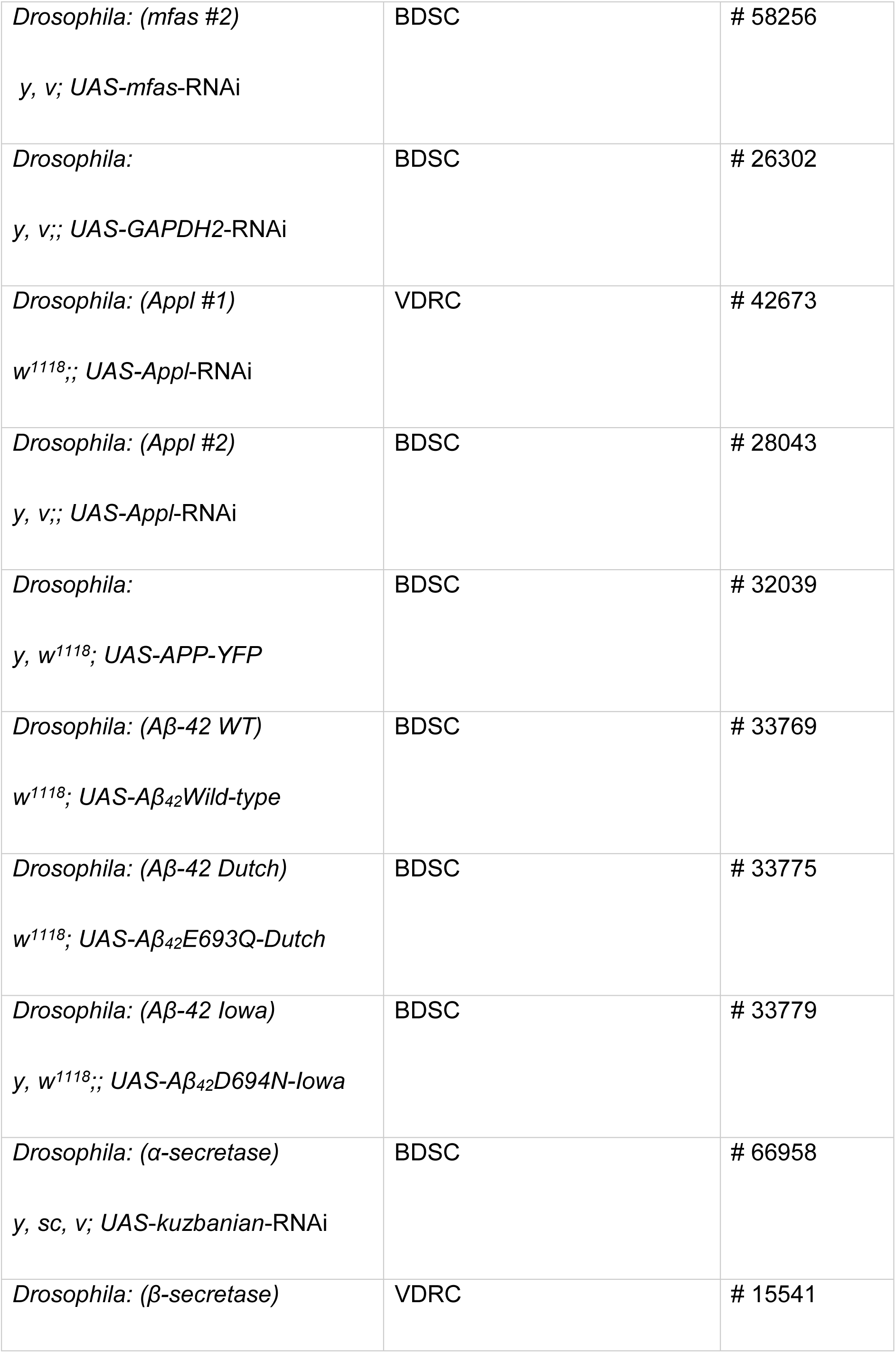
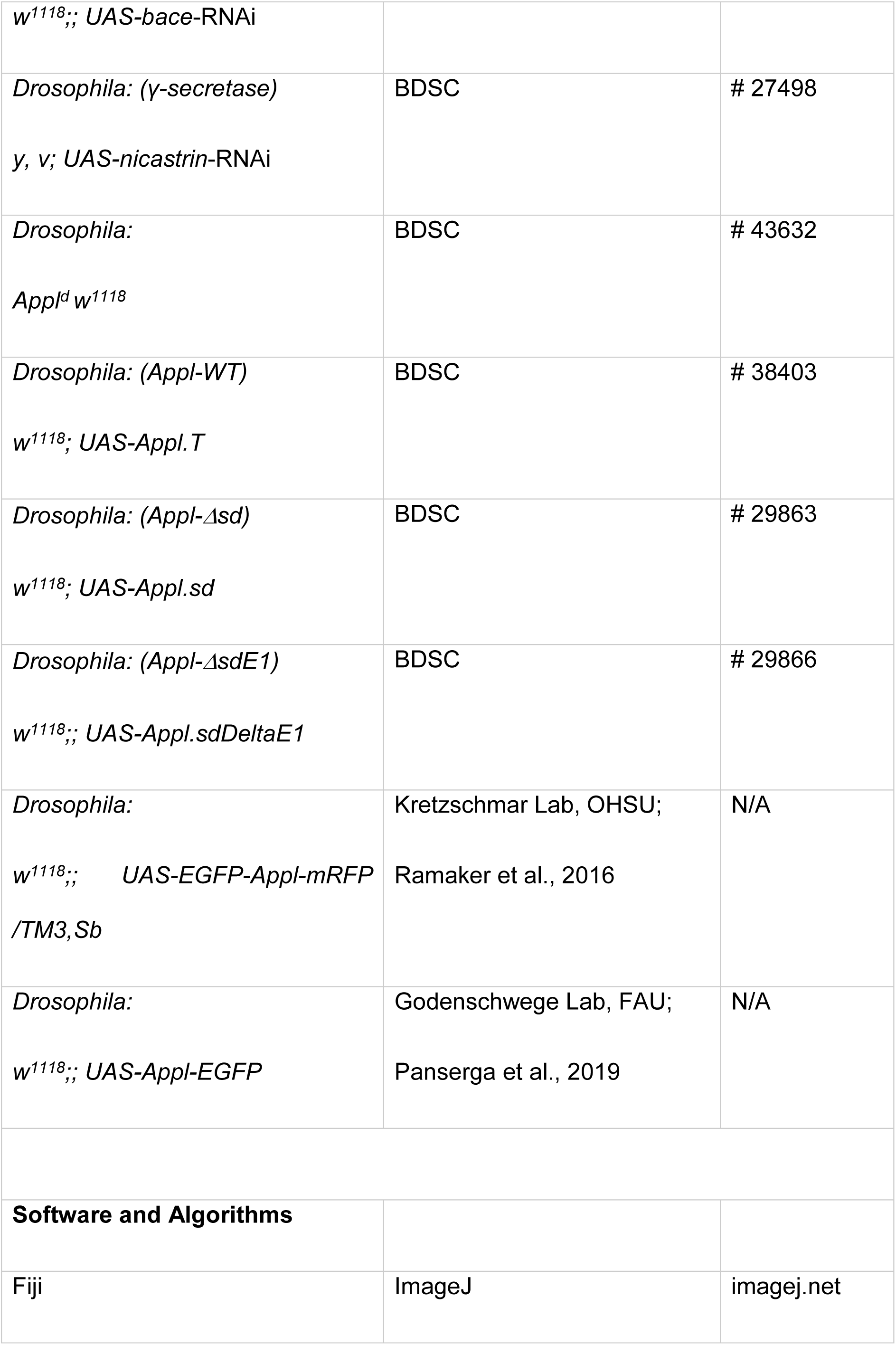
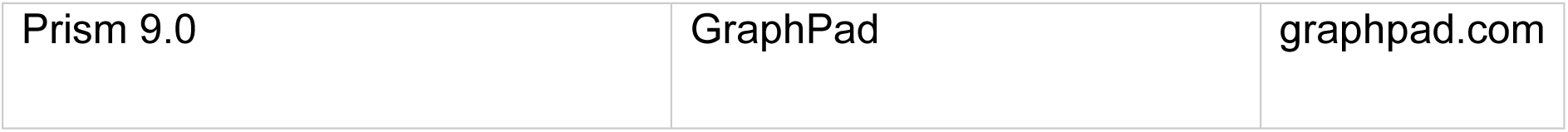

### Experimental model and subject details

#### Fly stocks and husbandry

All *Drosophila* strains used in this study are detailed in the key resource table. Where possible, RNAi lines were selected that have already been employed for gene knockdown in other studies, which are highlighted. The transgenic lines were acquired from Bloomington *Drosophila* Stock Centre (BDSC) and Vienna *Drosophila* Resource Centre (VDRC), unless otherwise stated: *GFP-mfas* gene trap (MI11275-GFSTF.2; BDSC 63204; Nagarkar-Jaiswal et al., 2015), *UAS-rosy*-RNAi (TRiP.HMS02827; BDSC 44106; Marie et al., 2023), *UAS-mfas*-RNAi #1 (TRiP.HMC03645; BDSC 52905; Ni et al., 2011) and #2 (TRiP.HMJ22320; BDSC 58256; Ni et al., 2011), *UAS-GAPDH2*-RNAi (TRiP.JF02072; BDSC #26302; Spannl et al., 2020; Dar et al., 2021), *UAS-Appl-*RNAi #1 (GD3170; VDRC 42673; Goguel et al., 2011) and #2 (TRiP.JF02878; BDSC 28043; Singh and Mlodzik, 2012), *UAS-APP-YFP* (BDSC 32039; Gunawardena et al., 2003), *UAS-Aβ-42 WT* (BDSC 33769; Wu et al., 2017), *UAS-Aβ-42 Dutch* (BDSC 33775; Vitruvean), *UAS-Aβ-42 Iowa* (BDSC 33779; Vitruvean; Metsla et al., 2022), *UAS-α-secretase*-RNAi (TRiP.HMS05424; BDSC 66958; Tian et al., 2023), *UAS-β-secretase*-RNAi (GD5366; VDRC 15541; Bolkan et al., 2012), *UAS-γ-secretase*-RNAi (TRiP.JF02648; BDSC 27498; Restrepo et al., 2022), *Appl^d^* (BDSC 43632; Luo et al., 1992), *UAS-Appl-WT* (BDSC 38403; Torroja et al., 1999), *UAS-Appl-Δsd* (BDSC 29863; Torroja et al., 1999), *UAS-Appl-ΔsdE1* (BDSC 29866; Torroja et al., 1999), *w^1118^* (provided by L. Partridge, UCL, UK), *UAS-GFP-Appl-RFP* (provided by D. Kretzschmar, OHSU, USA; Ramaker et al., 2016), *UAS-Appl-GFP* (provided by T. Godenschwege, FAU, USA; Panserga et al., 2019), *CFP-Rab6* and *YFP-Rab11* fusion genes at endogenous *Rab* locus (provided by S. Eaton, Max Plank, Germany). SC-specific temperature-sensitive driver lines were generated by combining *dsx-GAL4* (provided by S. Goodwin, Oxford, UK; Rideout et al., 2013) with ubiquitously expressed repressor *tub-GAL80^ts^* (BDSC 7108) to produce *tub-GAL80^ts^; dsx-GAL4* (Corrigan et al., 2014). This driver line was additionally combined with endogenously fluorescent tagged *GFP-mfas*, *CFP-Rab6* or *YFP-Rab11* to generate *CFP-Rab6, tub-GAL80^ts^; dsx-GAL4* (Wells et al., 2023), *tub-GAL80^ts^; YFP-Rab11, dsx-GAL4* (Redhai et al., 2016), and *tub-GAL80^ts^; dsx-GAL4, GFP-mfas* (this study).

Flies were maintained at 25^°^C under a 12-hour light/dark cycle on standard cornmeal agar medium [12.5 g agar (F.Gutlind & Co. Ltd), 75 g cornmeal (B. T. P. Drewitt), 93 g glucose (Sigma-Aldrich, #G7021), 31.5 g inactivated yeast (Fermipan Red, Lallemand Baking), 8.6 g potassium sodium tartrate tetrahydrate (Sigma-Aldrich, #S2377), 0.7 g calcium chloride dihyrdrate (Sigma-Aldrich, #21907), and 2.5 g nipagin (Sigma Aldrich, #H5501) dissolved in 12 ml ethanol, per litre]. They were transferred onto fresh food every 3 - 4 days. Female flies carrying the driver line *tub-GAL80^ts^; dsx-GAL4* alone or in combination with *GFP-mfas*, *CFP-Rab6* or *YFP-Rab11* were crossed with male flies carrying UAS-transgenes to induce a temperature-controlled SC-specific expression of target genes. These crosses were maintained at 25^°^C. Virgin male offspring were collected upon eclosion and typically transferred to 29^°^C for six days to activate post-developmental SC-specific transgene expression. Dissection and imaging were then performed.

### Method details

#### Preparation of Accessory glands for imaging

Accessory glands were dissected and prepared as described in Fan et al., 2020 and Wells et al., 2023. For live-cell imaging, six-day-old adult male virgin flies were anaesthetised using CO_2_. These flies were submerged in ice-cold 1X PBS (Thermo Fisher Scientific) during the micro-dissection procedure. The male reproductive tract was pulled out of the body cavity by carefully tweezing the last abdominal segment. The testes, seminal vesicles, ejaculatory bulb, fat tissues and the gut were gently removed to avoid tissue folding and interference during accessory gland imaging. The glands were then incubated with 500 nM Lysotracker Red (Thermo Fisher Scientific) for 5 minutes on ice, followed by a wash with ice-cold 1x PBS. The *ex vivo*-prepared glands were stably mounted between two coverslips (rectangular: Coverslip No.1, 22 mm x 50 mm, Fisher, #1237-3128, and round: coverslip No.1, 13 mm, #49492, VWR) in a drop of 1x PBS, held together by a custom-built metal holder. Excess PBS was removed using a filter paper until the glands were slightly flattened and ready for imaging.

For confocal analysis, micro-dissections were performed in PBS containing 4% paraformaldehyde (Sigma-Aldrich). The glands were fixed for 20 minutes at room temperature and then washed in 1x PBS for 3x 5 minutes prior to mounting onto SuperFrost microscope slides (VWR). After removing excess PBS using Whatman paper (Whatman), the glands were immersed in a drop of Vectashield with DAPI (Vector Laboratories) and stably positioned with a cover slip (22 mm x 22 mm, 0.13-0.17mm, Fisher).

#### Live-cell imaging, deconvolution and time-lapse movies

Based on Wells et al., 2023, live SCs were imaged at room temperature using a DeltaVision Elite wide-field fluorescence deconvolution microscope (GE Healthcare Life Sciences) at 100x (Olympus UPlanSApo NA 1.4; oil objective), using immersion oil with refractive index of 1.514 (Cargille labs) and a manual auxiliary magnification of 1.6x. An EMCCD Evolve-512 camera was used to capture images. Three SCs were imaged per accessory gland for ≥10 individual virgin males. The images acquired were typically z-stacks spanning a depth of 8-12 μm with a z-distance of 0.2 μm. The Resolve 3D-constrained iterative deconvolution algorithm within SoftWoRx 5.5 Software (GE Healthcare Life Sciences) was subsequently used to deconvolve z-stack images to improve the image quality prior to analysis.

For time-lapse imaging experiments, samples were prepared using the same method except that incubation with Lysotracker Red (Thermo Fisher Scientific) was typically excluded unless compartment acidification was being assessed. Time-lapse imaging was conducted using the DeltaVision Elite wide-field fluorescence deconvolution microscope (GE Healthcare Life Sciences) using identical methods to those described above with the following exceptions. Firstly, auxillary magnification was not utilised, in order to reduce the effects of cell/tissue drift over time, except for the control, *rosy-RNAi* cell in Movie S1. Secondly, whilst z-stacks of entire cells were acquired for time-lapse imaging experiments, for all but Movie S2, the z-distance between individual slices was increased from 0.2 μm to between 0.5 μm and 0.8 μm to limit photobleaching and phototoxicity. The gap between timepoints during these experiments was 60 seconds for all videos shown here except Movie S2, where the time interval was 90 seconds; additional videos were acquired with gaps of up to 120 seconds between frames. Finally, deconvolution was used as described above, but the final results of deconvolution were saved as 32-bit floating point images.

In order to best show all structures and fluorescence in compartments of interest, the images displayed in videos were made using z-projections that included the fluorescent signal from every z-slice within the bounds of the relevant compartment, typically four to eight image slices in the GFP-MFAS channel. For the DIC channel, a single representative z-slice was used generally taken from the centre-most point of the compartment.

#### Fixed accessory glands imaging

Fixed samples were imaged on a Zeiss LSM980 with Airyscan 2 Super-resolution upright laser scanning confocal microscope equipped with 10x (Zeiss 0.45 NA; dry) and 40x (Zeiss 1.30 NA; oil; Zeiss immersion oil, refractive index 1.518) objectives. High resolution images of the accessory gland lumen were acquired using the 40x objective with 0.7x zoom on the ZEN blue suite Software (Zeiss).

### Analysis and parameters

#### DCG phenotypes and number of mature compartments

Deconvolved images were analysed using Fiji/ImageJ. Number of intact DCGs marked by GFP-MFAS and DCG-containing compartments were scored using StarDist 2D plugin, DSB 2018 model which identifies each core as a separate entity. In the case of *mfas* knockdowns, the number of compartments were quantified manually using the Differential Interference Contrast (DIC) channel.

Abnormal cores included both those having a GFP-negative (GFP^-^) centre and those with mini-/deformed cores, with the exception of cells overexpressing the *Appl^sdΔE1^* mutant, which exhibited an additional peripheral network-like compartment phenotype. DCGs were scored as containing a GFP^-^ centre if they contained a non-fluorescent centre with ≥ 1 µm diameter, while the mini-/deformed core phenotype was defined by the presence of multiple small cores of diameter ≥ 0.5 µm and/or a misshapen core, where the ratio of the lengths of the longest and shortest DCG axes was > 1.4. Compartments with the peripheral network-like phenotype failed to form a round central DCG and most of the aggregated GFP-MFAS was in close proximity to the limiting membrane. These latter abnormal cores/compartments were manually scored using the z-stack in both DIC and Merge channels. The percentage abnormal and mini-/deformed cores was calculated relative to the total number of non-acidic DCG- (GFP-MFAS-) containing compartments per SC.

Intact or abnormally shaped DCGs that maintained contact with the limiting membrane through several planes in the z-stack were scored manually and were represented as a percentage relative to the total number of DCG compartments per SC.

To determine the frequency at which mini-cores either collide or overlap along the same z-axis in individual compartments, a single mini-core associated with the limiting membrane of each compartment was selected. Each instance of the mini-core’s fluorescent signal moving to overlap with other mini-cores and/or being in contact with another mini-core at the start of the movie was considered as a mini-core overlap. Images were acquired each minute over a duration of 30 minutes and the total number of overlaps was recorded.

To quantify compartments marked by either the CFP-Rab6 or YFP-Rab11 fusion protein expressed from the endogenous *Rab* locus, fluorescently labelled compartments were manually examined using z-stacks for the CFP or YFP channels (Wells et al., 2023). The proportion of these compartments containing *Rab6/11*-positive exosomes was established by evaluating which compartments contained internal fluorescent puncta in these genetic backgrounds.

#### Acidification of secretory compartments

Mature DCG compartments that are associated with acidic lysosomal structures and potentially undergoing lysosomal clearance (the DCG acidification phenotype) were scored as acidified compartments. These compartments manifest different phenotypes depending on the stage of lysosomal clearance. Some have a single peripheral lysosomal structure with a slightly diffuse DCG that has maintained its shape, while others have completely diffuse GFP-MFAS and no obvious DCG structure. Some feature multiple lysosomal structures arranged in an arc around the compartment boundary, with or without diffuse GFP-MFAS. Others are >80% covered by acidified domains, but the compartments still retain their circular shape. The percentage of acidified compartments per SC was determined as a proportion of the total number of both non-acidic DCG-containing compartments and acidified compartments per SC.

To calculate the lysosomal area, freehand tool on Fiji was employed to outline and measure the area of the SC. Threshold and analyze particles tools were utilised to determine the lysosomal area in the complete projection of the lysotracker (RFP) channel. The percentage of lysosomal area per SC was then calculated relative to the total area of the SC.

#### GFP-MFAS-containing main cells

The Rectangle tool on Fiji was employed to determine the area of the field of view, and the freehand tool was employed to outline and measure the area of the SC. Subsequently, the threshold and analyse particles tools were applied to measure the area covered by GFP-MFAS in the main cells using the complete projection of the GFP channel. The percentage of main cell area containing GFP-MFAS was calculated within the field of view, excluding the SC area.

#### GFP-MFAS secretion into accessory gland lumen

The central region of fixed accessory glands was imaged to examine the lumenal phenotype. Using the rectangle tool, two squares of equal sizes (150 x 150 pixels) were marked in the middle of each arm, resulting in a total of four squares per animal. The mean gray value of these four areas was calculated using the same settings for all images, and presented as an average GFP-MFAS intensity per gland.

#### Image processing and preparation for figures

Live-cell images for this study were prepared using deconvoluted stacks collected from the DV Elite microscope. In all figures, the DIC channel showcased a single-slice image chosen for its optimal representation of SC morphology. Max-projections of slices ranging from 3-10 μm was employed for all fluorescent channels, specifically highlighting DCG compartments and Rab6-/Rab11-compartments with ILVs. The fluorescence intensity that best captured the phenotype was selected and utilised for image processing across all the live-cell data. Consistent projection of slices and settings were applied to generate images for each genotype shown in the figures. All images were cropped to identical dimensions, focusing primarily on SC details. To enhance contrast, sharpen tool in ImageJ was uniformly applied to all images ensuring unbiased and accurate data representation.

For fixed-gland images, a single slice that most accurately represented the phenotype was acquired using the LSM980 confocal microscope. To ensure consistency in fluorescent intensity quantification, the GFP gain settings were kept constant across all genotypes.

#### Human sEV isolation

Human HCT116 Rab11a-exosome analysis was undertaken using the techniques described in Fan et al., 2020 and Marie et al., 2023. Approximately 8–9 x 10^6^ HCT116 cells were seeded per 15 cm cell culture plate, using 10–15 plates per condition, and allowed to settle for 16–18 h in complete medium before the 24-h EV collection. This involved culture for 24 h in serum-free DMEM/F12 medium without L-glutamine (#21331046; Life Technologies), which was supplemented with 1% ITS (Insulin-Transferrin-Selenium; #41400045 Life Technologies), and either 2.00 mM or 0.15 mM glutamine (Life Technologies). Cells grew to approximately 90% confluence by the end of the collection period. The culture medium was centrifuged at 500 x *g* for 10 min at 4°C and 2000 x *g* for 10 min at 4°C to remove cells, debris and large vesicles. The supernatant was filtered to remove remaining large EVs using 0.22 μm filters (Mile).

The filtrate was concentrated to a volume of approximately 30 ml using a tangential flow filtration (TFF) unit with a 100 kDa membrane (Vivaflow 50R, Sartorius) coupled to a 230 V pump (Masterflex). The sEV suspension was then further concentrated using 100 kDa Amicon filters by sequential centrifugation at 4000 x g for 10 min at 4°C to give a final volume of 1 ml. This was injected into a size-exclusion column (column size 24 cm x 1 cm) containing Sepharose 4B (84 nm pore size) using an AKTA start system (GE Healthcare Life Science) and eluted with PBS, collecting 30x 1 ml fractions. Fractions corresponding to the initial “EV peak” (typically fractions two to five) were pooled in 100 kDa Amicon tubes to a final volume of approximately 100 μl.

#### Nanoparticle tracker analysis (NanoSight^®^) of sEVs

Using the NS500 NanoSight^®^, between three and five thirty-second videos were captured per EV sample at known dilution (normalised to protein mass of secreting cells). Particle concentrations were measured within the linear range of the NS500 (approximately 2–10 x 10^8^ particles per ml). Particle movement was analysed by NTA software 2.3 (NanoSight Ltd.) to determine particle size distribution and concentration.

#### Western analysis

Both cell lysates and EV preparations, which were lysed in RIPA or 1X sample buffer. Protein preparations were dissolved in either reducing (with 5% β-mercaptoethanol) or non-reducing (for CD63 and CD81 detection) sample buffer and heated to 90– 100°C for 10 min before loading. A pre-stained protein ladder (Bio-Rad) was also used. Proteins were separated by electrophoresis using 10% mini-PROTEAN precast gels (BioRad). For gels of sEV proteins, lanes were loaded with EV lysates extracted from the same protein mass of secreting cells. This ensured that changes in band intensity on the blots with glutamine depletion reflected a net change in secretion of the marker on a per cell basis (see Fan et al., 2020).

Proteins were wet-transferred to polyvinylidene difluoride (PVDF) membranes at 100 V for 1 h using a Mini Trans-Blot Cell (Bio-Rad). Membranes were blocked with either 5% milk (CD63 detection) or 5% BSA in TBS buffer with Tween-20 (TBST) for 30 min and probed overnight at 4°C with primary antibody diluted in blocking buffer. The membranes were washed for 3x 10 min with TBST, then probed with the appropriate secondary antibodies for 1 h at 22°C and then washed for 3x 10 min. Signals were detected using the enhanced chemiluminescent detection reagent (Clarity, BioRad) and a Touch Imaging System (BioRad). Relative band intensities were quantified by ImageJ.

Antibody suppliers, catalogue numbers and concentrations used were: mouse anti-Tubulin (Sigma #T8328, 1:4000), mouse anti-CD81 (Santa Cruz #23962, 1:500), mouse anti-CD63 (BD Biosciences # 556019, 1:500), rabbit anti-Syntenin-1 antibody (Abcam ab133267, 1:500), rabbit anti-Tsg101 (Abcam ab125011, 1:500), anti-GAPDH (DSHB hGAPDH-2G7, 1:250), mouse anti-Rab11 (BD Biosciences #610657, 1:500), anti-mouse IgG (H+L) HRP conjugate (Promega #W4021, 1:10000), anti-rabbit IgG (H+L) HRP conjugate (Promega #W4011, 1:10000).

#### Comparative proteomics analysis of HeLa cell sEV preparations

Four paired samples of conditioned medium (serum-free basal medium [DMEM/F12] supplemented with 1% ITS [Insulin-Transferrin-Selenium; #41400045 Life Technologies]) were collected over a 24-h period from glutamine-depleted and glutamine-replete HeLa cells, using the same culture conditions as for HCT116 cells, except that the cells were seeded at 4 x 10^6^ cells per plate. sEVs were isolated by size-exclusion chromatography, as described above.

Each sEV sample was then lysed in 100 μL RIPA buffer for 30 min on ice, followed by 10 min centrifugation at 17,000 x g at 4°C. The clear supernatants were transferred to fresh tubes. Proteins were reduced with DTT (final concentration 5 mM) and alkylated with Iodoacetamide (final concentration 20 mM) for 30 min each and then precipitated with methanol/chloroform. They were then mixed with 600 μl methanol, 150 μl chloroform and 450 μl ddH2O and centrifuged again for 3 min at 17,000 x g. The upper phase was carefully removed and a further 450 μl methanol added. After another centrifugation step for 5 min, supernatants were removed and discarded. The protein pellets were resuspended in 50 μl 100 mM TEAB and digested with 200 ng trypsin (Promega sequencing grade) overnight at 37°C.

Sample amounts were normalized based on protein concentration measurements for TMT labelling. Approximately 5.6 μg of peptides in 50 μl 100 mM TEAB were labelled with 0.06 mg of a TMT10plex label for 1 h, then quenched with 5 μl 400 mM Tris-HCl for 15 min. All eight samples were combined, desalted on SOLA HRP SPE cartridges (Thermo Scientific) and dried down in a vacuum centrifuge. Samples were resuspended in 2 % acetonitrile with 0.1% formic acid.

Samples were analysed on the Dionex Ultimate 3000/Orbitrap Fusion Lumos platform. Peptides were separated on a 50 cm, 75 μm ID EasySpray column (ES803; Thermo Fisher) on a 60-minute gradient of 2 to 35% acetonitrile (containing 0.1% formic acid and 5% DMSO) at flow rate of 250 nl/min. Data were acquired using the MultiNotch MS3 method, as described previously (McAlister et al., 2014).

Mass spectrometry raw data were analysed in Proteome Discoverer 2.1. Proteins were identified with Sequest HT against the UPR Homo sapiens database (retrieved February 2017). Mass tolerances were set to 10 ppm for precursor and 0.5 Da fragment mass tolerance. TMT10plex (N-term, K), Oxidation (M) and Deamidation (N, Q) were set as dynamic modifications, alkylation (C) as a static modification. The mass spectrometry proteomics data have been deposited to the ProteomeXchange Consortium via the PRIDE (Vizcaíno et al., 2016) partner repository with the dataset identifier TBC???.

### Statistical analysis

For comparing multiple experimental genotypes with the control, we applied the non-parametric Kruskal-Wallis test followed by Dunn’s multiple comparisons post-hoc test. When comparing two groups, a non-parametric Mann-Whitney test was used. These statistical analyses were performed on GraphPad Prism. All graphs displayed in the figures show the mean value for each genotype and include error bars representing standard error of mean (SEM). Each ‘n’ in the analysis corresponds to the number of SCs analysed for each genotype, assessed using at least 10 independent AG lobes.

For Western blots, relative signal intensities were analysed using the Kruskal–Wallis test.

## Acknowledgements

We are grateful to all the staff at the Micron Bioimaging Facility for their support, to Amy Cording for her technical assistance, and to Rapheal Heilig and Benedikt Kessler for their support with the proteomics analysis. We thank S. Eaton, S. Goodwin, E. Prince, F. Karch, L. Partridge, T. Godenschwege and D. Kretzschmar, as well as the Bloomington and Vienna *Drosophila* Stock Centres for *Drosophila* stocks. We acknowledge the support of the BBSRC (BB/L007096/1, BB/N016300/1, BB/R004862/1, BB/W00707X/1, BB/W015455/1), Cancer Research UK (C19591/A19076, C602/A18974) and the National Science and Technology Council, Taiwan (111-2320-B-008-001-MY2) to S.J.F. P.J.S. was the recipient of a grant from the Balliol Interdisciplinary Institute (BII), a subsidiary of Balliol College, Oxford. A.W. was supported by a Krebs Memorial Scholarship from the Biochemical Society, and R.F. by Wellcome (097813/11/Z) and the John Fell Fund (133/075). For the purpose of Open Access, the author has applied a CC BY public copyright licence to any Author Accepted Manuscript (AAM) version arising from this submission.

## Supplementary Information

### Supplementary Figures

**Figure S1.**
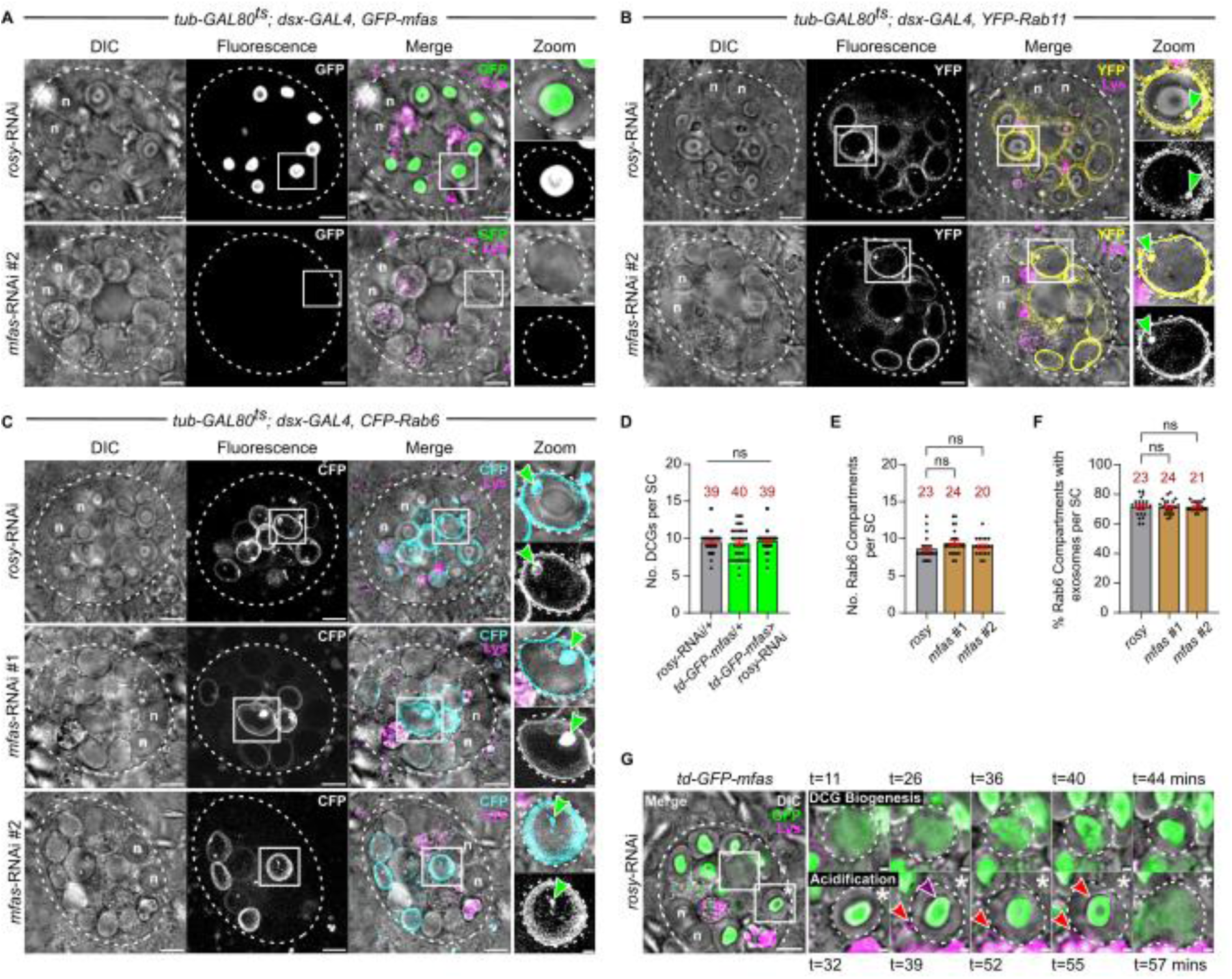
*Drosophila* TGFBI selectively drives DCG assembly in SCs, related to Figure 1. (A-C) *Ex vivo*, wide-field fluorescence micrographs and DIC images of SCs from 6-day-old males expressing SC-specific *rosy*-RNAi or either of two independent *mfas*- RNAis with the *GFP-mfas* gene trap (A), *YFP-Rab11* (B) or *CFP-Rab6* (C). Zoom panels show absence of DCGs following *mfas* knockdown, but presence of Rab-positive ILVs (green arrowheads) in compartments. (D) Expressing *rosy*-RNAi in adult SCs using the *tub-GAL80^ts^; dsx-GAL4, GFP-mfas* SC-specific GAL4 driver line has no effect on DCG compartment number compared to controls. (E and F) Knockdown of *mfas* with two independent RNAis has no effect on numbers of Rab6-positive large compartments (E) or the proportion of these compartments containing Rab6-positive ILVs (F). (G) Stills from time-lapse movie of DCG biogenesis and DCG acidification in SC from 6-day-old male expressing *GFP-mfas* gene trap. For compartment marked by white box (top row of zoomed images), a single DCG forms rapidly from a GFP-MFAS cloud. For compartment marked by white box and asterisk (bottom row), small LysoTracker Red-positive compartments (red arrowheads) contact and spread around the periphery of the DCG compartment, then start to acidify the lumen around the DCG (66 mins) before rapid dispersion of the core (67.5 mins). Purple arrowhead marks DCG that can persist for twenty minutes following the start of the acidification process. *td-GFP-mfas* = *tub-GAL80^ts^/+; dsx-GAL4, GFP-mfas/+*. In all images, approximate cell boundary and compartment boundaries are marked with dashed white line; n = nuclei of binucleate cells; LysoTracker Red (magenta) marks acidic compartments. Scale bars = 5 µm and 1 µm in Zoom. For bar charts, data were analysed using the Kruskal-Wallis test, followed by Dunn’s multiple comparisons post hoc test; n = number above bar, ns = not significant.

**Figure S2.**
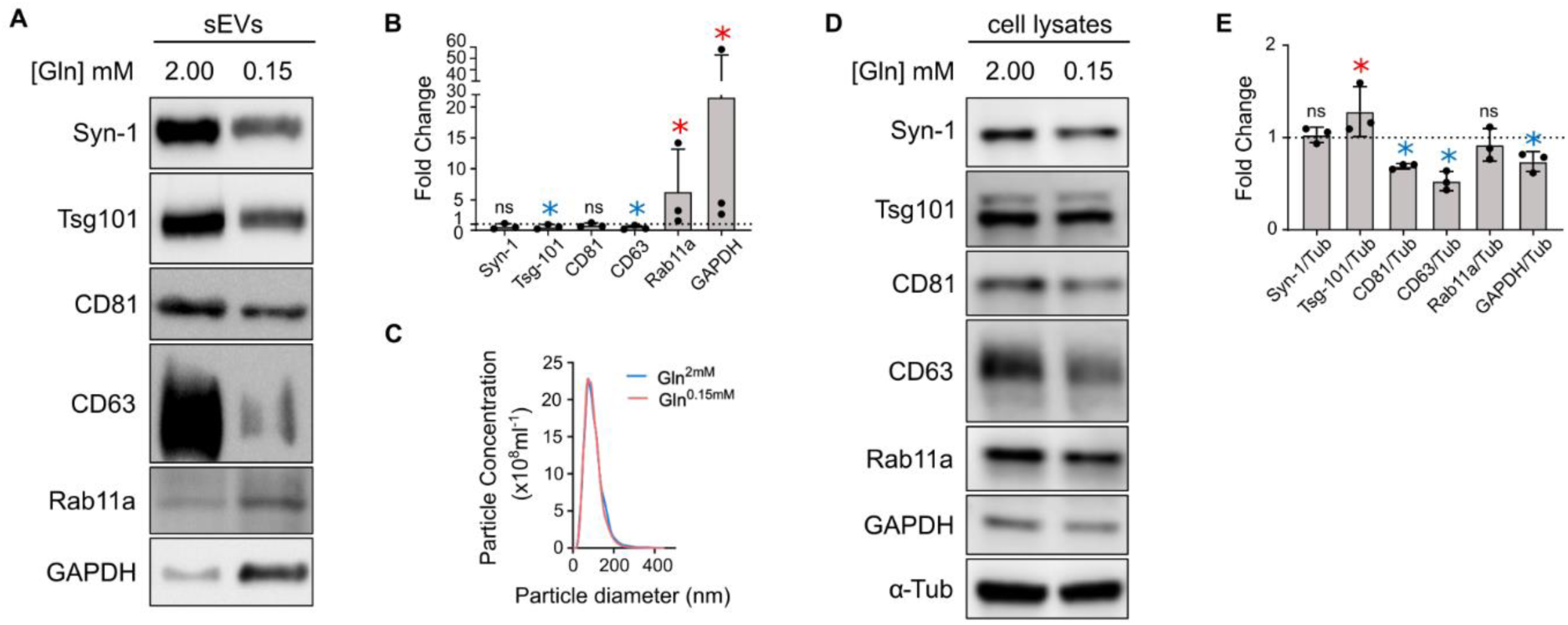
GAPDH is enriched in human HCT116 Rab11a-exosome preparations, related to Figure 2. (A and B) Western blot analysis of putative exosome markers in sEV preparations concentrated by ultracentrifugation and collected from HCT116 cells cultured under glutamine-replete (2.00 mM) and glutamine-depleted (0.15 mM) conditions for 24 h (A). Loading is based on cell lysate protein levels, so that secretion is compared on a per cell basis. Syn-1 = Syntenin-1. Bar charts show relative change in exosome protein levels in sEVs collected under glutamine-depleted versus -replete conditions, ie. Rab11a-exosome-enriched versus -depleted sEV preparations respectively (B). (C) Nanosight Tracking Analysis of EV size and number for diluted sEV samples (normalised to cell lysate protein levels) from cells cultured in glutamine-replete and glutamine-depleted conditions for 24 h, as in Figure S2A. (D and E) Western blot analysis of putative exosome proteins in lysates from HCT116 cells cultured under glutamine-replete (2.00 mM) and glutamine-depleted (0.15 mM) conditions for 24 h (D). Equal amounts of protein were loaded. Bar chart shows relative abundance of putative exosome proteins in these two lysates, normalised to relative abundance of tubulin (E). The elevated levels of Rab11a and GAPDH in Rab11a-exosome-enriched sEV preparations cannot be explained by increased cellular expression.

**Figure S3.**
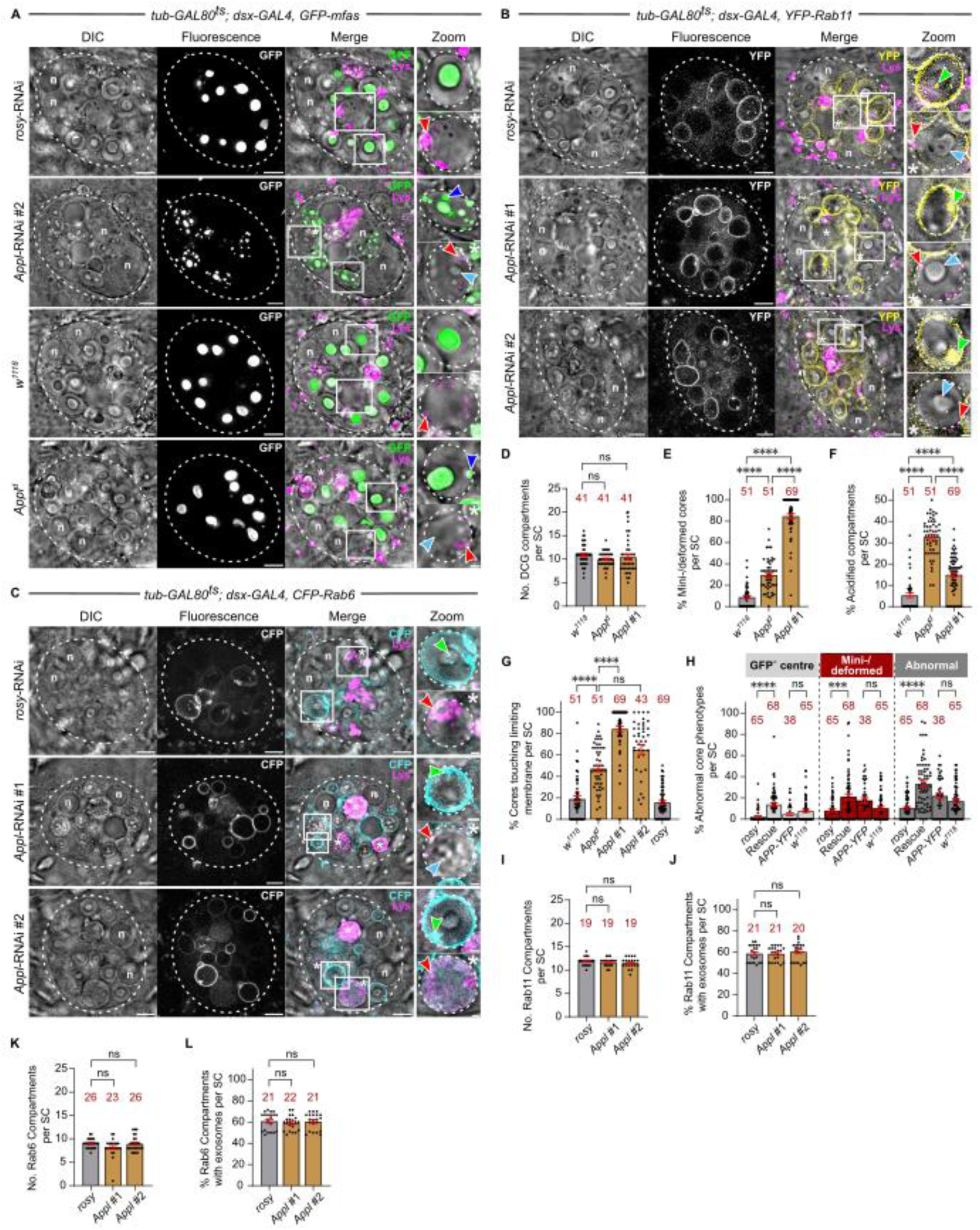
*Drosophila* APPL regulates formation of large DCGs in SCs, related to Figure 3. (A) *Ex vivo*, wide-field fluorescence micrographs and DIC images of SCs from 6-day-old males expressing *GFP-mfas* gene trap and SC-specific *rosy*-RNAi or *Appl*-RNAi #2 (top two rows), or without additional transgenes, but in a control *w^1118^* or *Appl^d^* mutant background (bottom two rows). Note that following knockdown of *Appl*, large secretory compartments contain multiple mini-cores (blue arrowheads), labelled with GFP-MFAS and visible by DIC (white box and upper Zoom panel). *Appl^d^* mutant SCs contain distorted DCGs that frequently contact the compartment’s limiting membrane and sporadic mini-cores (white box, top Zoom panel). The DCG acidification phenotype is also more commonly observed in *Appl* knockdown and *Appl* mutant backgrounds (white asterisks and white boxes marked by asterisks, shown in lower Zoom panels; red arrowheads mark acidic microdomains). DCGs that have not yet been dissipated marked with light blue arrowheads. (B and C) *Ex vivo*, wide-field fluorescence micrographs and DIC images of SCs from 6-day-old males expressing SC-specific *rosy*-RNAi or either of two independent *Appl*- RNAis with *YFP-Rab11* (B) or *CFP-Rab6*. Zoom panels show mini-core phenotype (white box, top panel) with intra-compartmental Rab puncta (green arrowheads), and DCG acidification phenotype with no Rab association (white box marked with asterisk, bottom panel). (D-G) Bar charts comparing the effects of the *Appl^d^* null mutant with SC-specific knockdown of *Appl* and controls. *Appl^d^* does not affect the number of DCG compartments (D). Although it typically does not induce mini-core formation, DCGs are often deformed (E) and touch the compartment’s limiting membrane (G). The DCG acidification phenotype is particularly prominent in *Appl^d^* mutant SCs (F). (H) Bar chart showing DCG phenotypes induced by APP-YFP expression with and without *Appl* knockdown versus control SCs. Note that in *Appl* knockdown cells rescued by APP-YFP, about 10% of DCGs are mis-assembled, lacking GFP-MFAS at their centre (GFP^-^ centre), perhaps because cleaved APP-YFP is involved in priming the aggregation of proteins at the centre of the compartment in this genetic background. (I-L) Bar charts showing that knockdown of *Appl* with two independent RNAis has no effect on numbers of Rab11-positive (I) or Rab6-positive (K) large compartments or on the proportion of these compartments containing Rab11-positive (J) or Rab6-positive (L) ILVs. In all images, approximate cell boundary and compartment boundaries are marked with dashed white line; n = nuclei of binucleate cells; LysoTracker Red (magenta) marks acidic compartments. Scale bars = 5 µm and 1 µm in Zoom. For bar charts, data were analysed using the Kruskal-Wallis test, followed by Dunn’s multiple comparisons post hoc test; n = number above bar, ***P<0.001, ****P<0.0001, ns = not significant.

**Figure S4.**
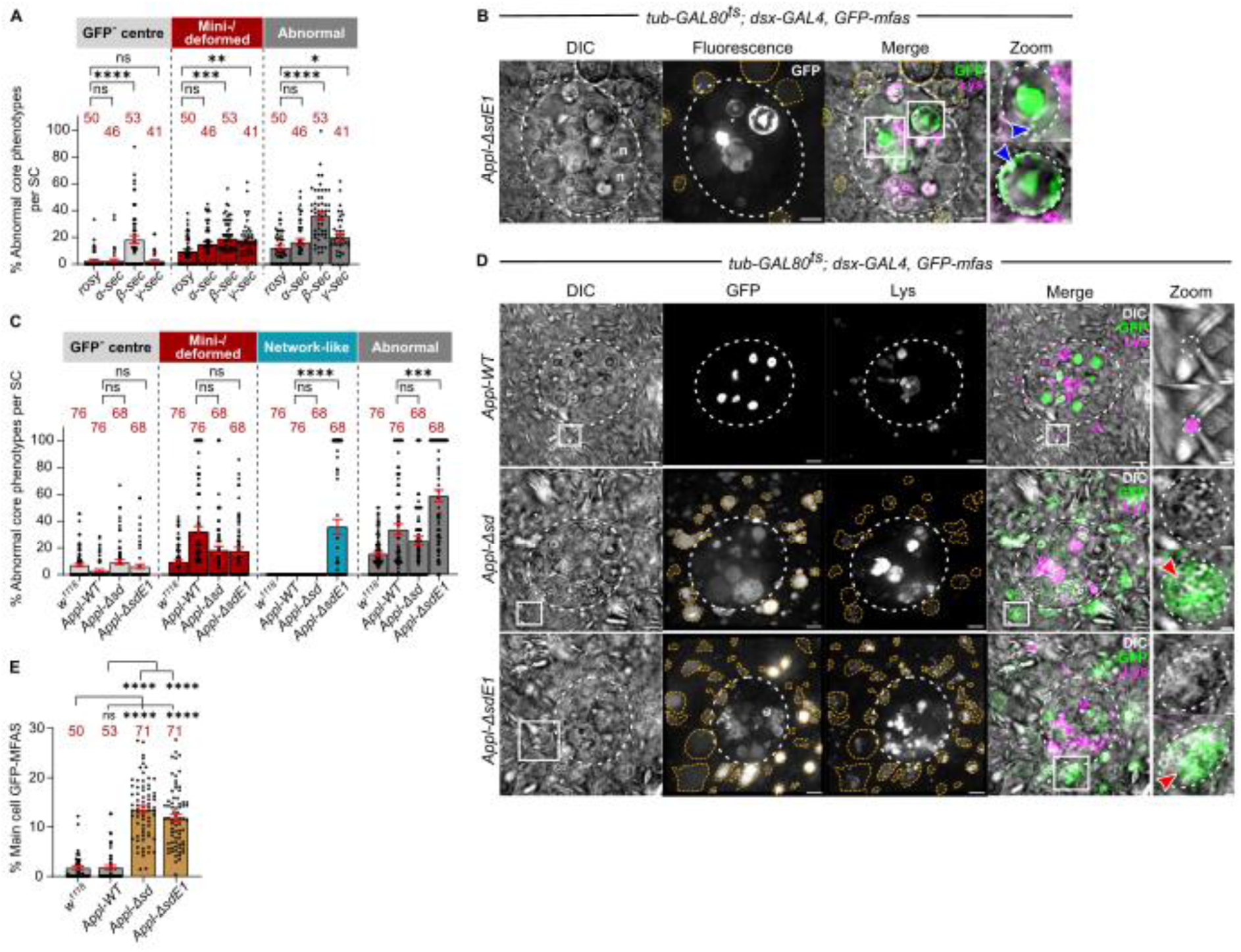
*Drosophila* APPL and its cleavage regulate normal DCG formation, and the uptake of GFP-MFAS by other cells, related to Figure 6. (A) Bar chart showing the different DCG phenotypes observed following knockdown of *α-*, *β-* and *γ-secretases* compared to *rosy* knockdown control. Note that *β-secretase* knockdown induces formation of DCGs that lack GFP-MFAS at their centre, while knockdown of both *β-* and *γ-secretases* produces increased numbers of deformed DCGs. (B) *Ex vivo*, wide-field fluorescence micrographs and DIC images of SC from 6-day-old male expressing *GFP-mfas* gene trap and APPL-ΔsdE1 (Figure 4B). Note two abnormal DCG compartments that have a central abnormally shaped DCG, but also contain peripheral GFP-MFAS aggregates (blue arrowheads in compartments outlined with white boxes shown in Zoom panels). (C) Bar chart showing that overexpression of APPL-WT, APPL-Δsd and APPL-ΔsdE1 produces abnormal DCGs, with APPL-ΔsdE1 generating a unique network phenotype. (D) *Ex vivo*, wide-field fluorescence micrographs and DIC images of SC and surrounding main cells from 6-day-old male expressing *GFP-mfas* gene trap and wild-type APPL (APPL-WT), APPL-Δsd or APPL-ΔsdE1. Note abnormal accumulation of GFP-MFAS in main cell compartments that typically exhibit limited LysoTracker Red staining. Highly enlarged, GFP-containing acidic compartments, which are formed in main cells when APPL-Δsd and APPL-ΔsdE1 are expressed in SCs, are marked by yellow arrowheads and one example is outlined by a white box and shown in Zoom panels (DIC alone and DIC/Merge; red arrowheads mark acidic microdomains). (E) Bar chart showing the accumulation of GFP-MFAS in main cells of 6-day-old males overexpressing SC-specific APPL-Δsd and APPL-ΔsdE1 versus APPL-WT and *w^1118^* controls that do not express an APPL transgene, expressed as percentage of total main cell area containing GFP. In all images, approximate SC boundary is marked with dashed white line; n = nuclei of binucleate cells; LysoTracker Red (magenta) marks acidic compartments in B and D. Scale bars = 5 µm and 1 µm in Zoom. For bar charts, data were analysed using the Kruskal-Wallis test, followed by Dunn’s multiple comparisons post hoc test; n = number above bar, *P<0.05, **P<0.01, ***P<0.001, ****P<0.0001, ns = not significant.

**Figure S5.**
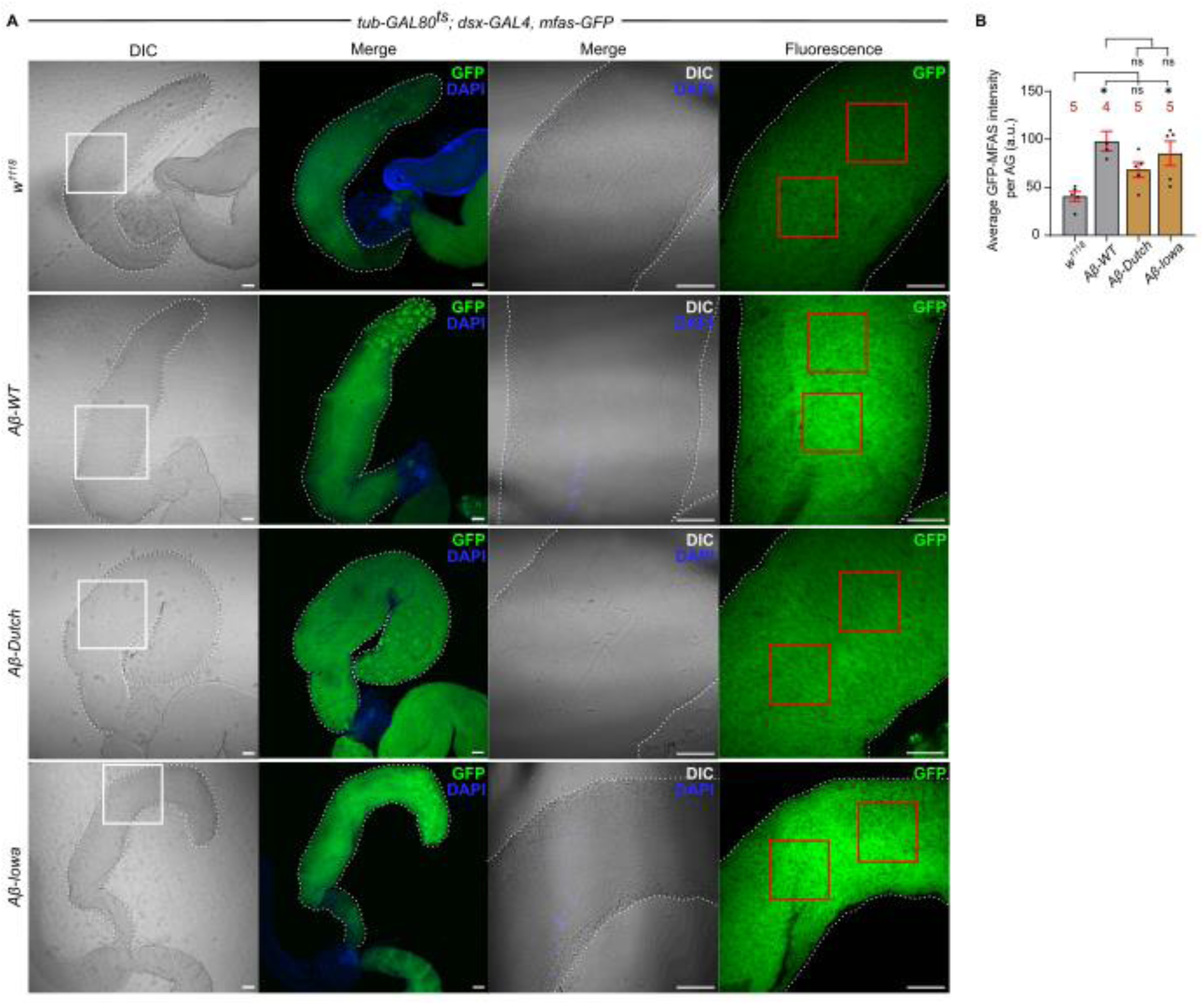
GFP-MFAS secretion and extracellular distribution is not noticeably affected by pathological Aβ-peptide expression in SCs, related to Figure 7. (A) Confocal micrographs (DIC and fluorescence) of fixed accessory glands of 6-day-old male expressing *GFP-mfas* gene trap and either no other transgene, or wild type Aβ-42 peptide, or either the Iowa or Dutch mutant Aβ-42 peptides. Two panels on right-hand side are magnified images of region in white box in left-hand panel. Two red boxes on right-hand panels mark two lumenal regions from which GFP signal was measured in these specific images. (B) Bar chart showing average GFP intensity in accessory gland lumen for the four genotypes shown. All glands were stained with DAPI to mark nuclei. Scale bars = 50 µm. For bar chart, data were analysed using the Kruskal-Wallis test, followed by Dunn’s multiple comparisons post hoc test; n = number above bar, *P<0.05, ns = not significant.

### Supplementary Table

**Table S1.**
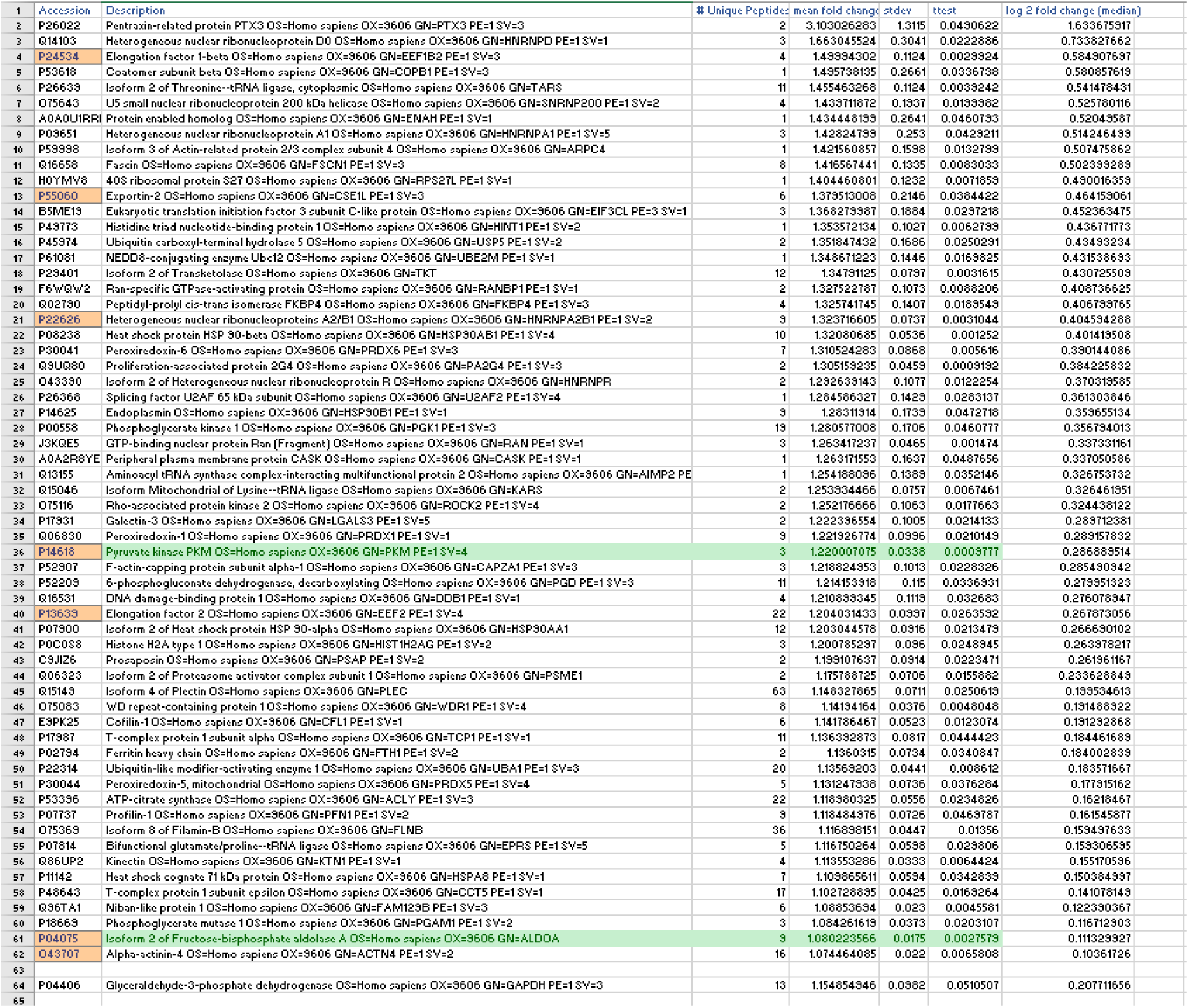
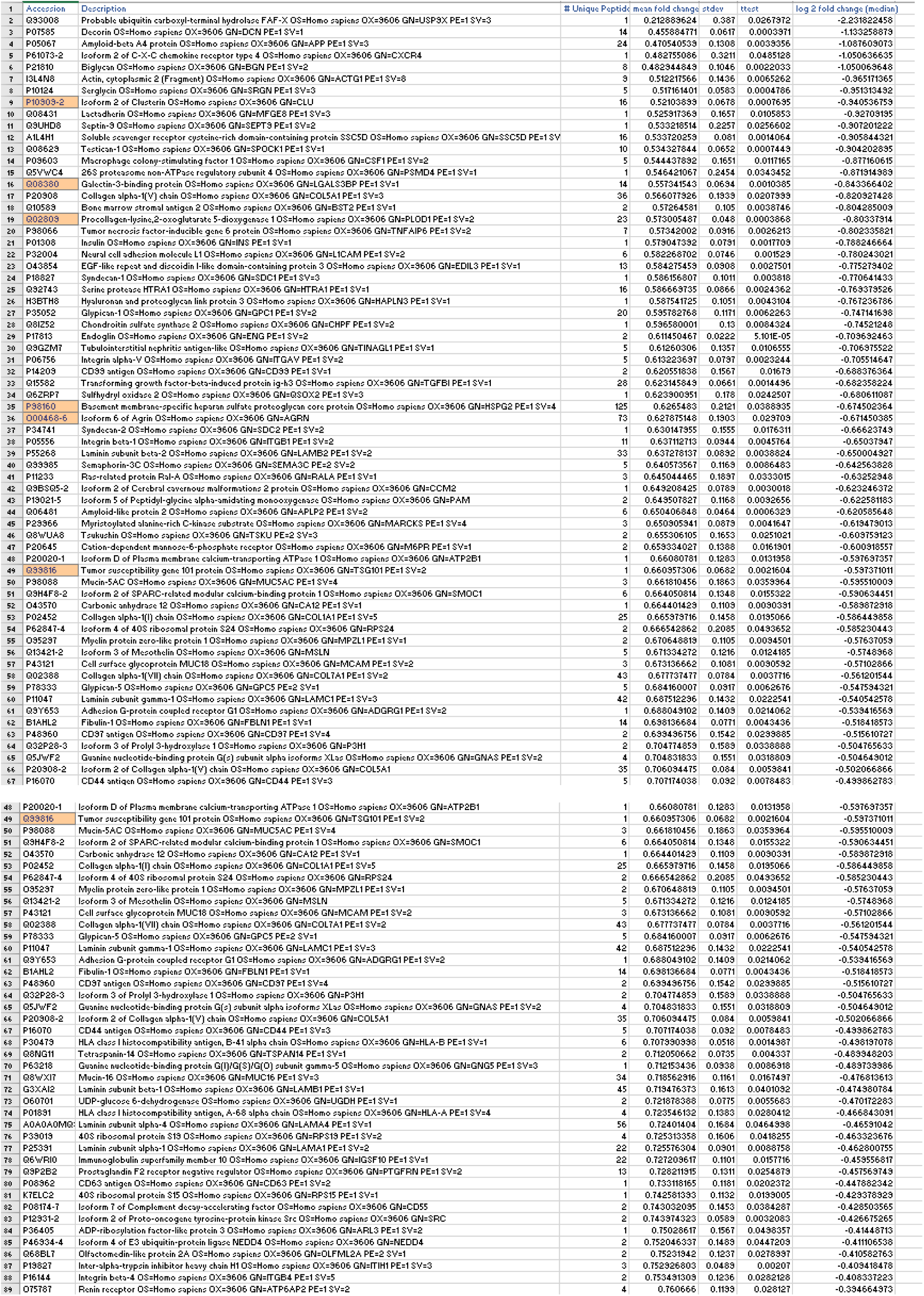
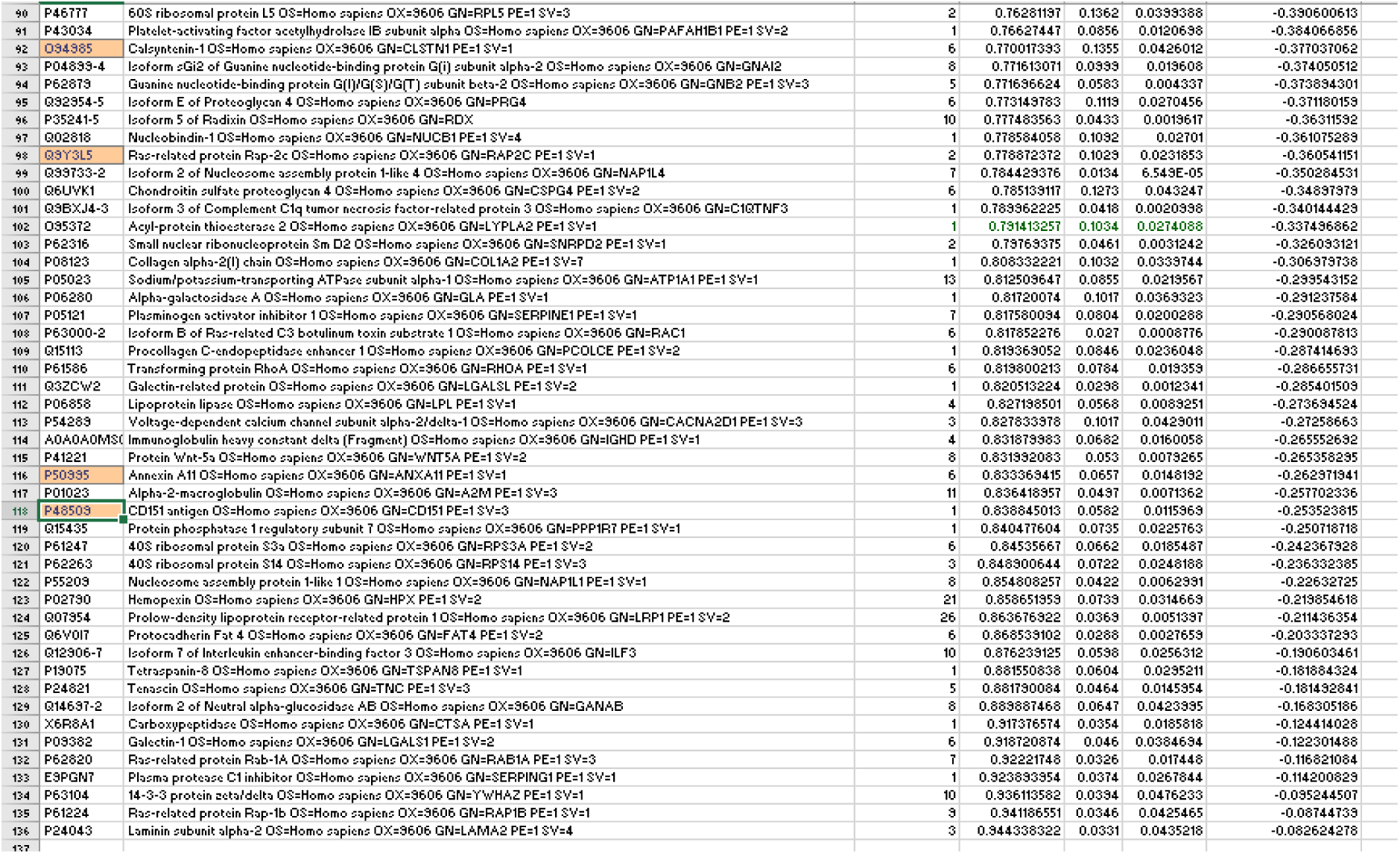
List of proteins increased or decreased in HeLa cell Rab11a-exosome preparations, related to Figure 2. Tables show lists of HeLa cell proteins with statistically significant increased or decreased levels in Rab11a-exosome-enriched sEV preparations versus sEV preparations collected under normal glutamine-replete (2 mM) culture conditions and concentrated by size-exclusion chromatography. Secretion of Rab11a-exosomes was induced by culturing in 0.02 mM glutamine. Data were generated from four pair-wise comparisons using the TMT labelling approach, so that all samples were analysed in the same MS run. Orange highlighted accession numbers denote those proteins that were similarly increased or decreased in a previous comparative TMT proteomics analysis of HCT116 sEVs collected under glutamine-depleted versus -replete conditions and concentrated by ultracentrifugation (Marie et al., 2023). Green highlighting denotes the two glycolytic enzymes, ALDOA and PKM.

### Supplementary Movies

**Movie S1** – **Time-lapse movie of SC DCG biogenesis in *GFP-mfas* genetic background**, related to Figure 1H.

**Movie S2** – **Time-lapse movie of SC DCG biogenesis in *GFP-mfas* genetic background**, related to Figure S1G.

**Movie S3** – **Time-lapse movie of SC DCG biogenesis in *GAPDH2* knockdown cell with *GFP-mfas* marker**, related to Figure 2E.

**Movie S4 – Time-lapse movie of SC DCG biogenesis in *Appl* knockdown cell with *GFP-mfas* marker**, related to Figure 3G.

**Movie S5** – **Time-lapse movie of DCG biogenesis in SC expressing Aβ-42 Dutch mutant in *GFP-mfas* genetic background**, related to Figure 7F.

**Movie S6** – **Time-lapse movie of DCG biogenesis in SC expressing Aβ-42 Iowa mutant in *GFP-mfas* genetic background**, related to Figure 7G.

